# Continuous DNA Methylation Deconvolution-Based Surrogate for B-Cell Differentiation State in CLL

**DOI:** 10.1101/2025.09.16.676535

**Authors:** Jeffrey Hage, Lauren Wainman, Fiyinfoluwa Kolawole, Fred Kolling, Pauline Patzke, Daniel Schoolcraft, Sarah Spracklin, Prabhjot Kaur, Brock Christensen

**Affiliations:** Department of Epidemiology, Geisel School of Medicine, Dartmouth College, Lebanon, NH, USA; Dartmouth Cancer Center, Dartmouth-Hitchcock Medical Center, Lebanon, NH, USA; Department of Pathology and Laboratory Medicine, Dartmouth-Hitchcock Medical Center, Geisel School of Medicine at Dartmouth, Lebanon, NH, USA; Department of Molecular and Systems Biology, Geisel School of Medicine, Dartmouth College, Lebanon, NH, USA

**Keywords:** CLL, Chronic Lymphocytic Leukemia, DNA Methylation, Deconvolution, Epigenetics, Cell-of-Origin, Transcription Factor, Tumor Burden, B-naive-like, B-memory-like

## Abstract

Chronic Lymphocytic Leukemia (CLL) is clinically divided into *IGHV* mutated (M-CLL) and *IGHV* unmutated (U-CLL) subtypes, which are thought to arise from distinct cells of origin along the B-cell differentiation pathway. We measured genome-scale DNA methylation in purified CLL samples (***n* = 89**) and utilized reference-based cell deconvolution techniques to develop a continuous metric of epigenetic similarity across a B-naive-like to B-memory-like scale (B-Index). B-Index accurately classifies CLL into clinical subtypes (98.8%), has a stronger epigenetic signal than *IGHV* gene percent identity, and demonstrates additional epigenetic signal within the M-CLL subgroup. We demonstrate that U-CLL is epigenetically more similar to B-memory than B-naive cells and reconcile previous reports of a B-naive-like epigenetic signal. The B-memory-like program of U-CLL is enriched for binding sites of transcription factors related to the germinal center activation pathway. Our findings provide epigenetic evidence for discerning CLL mechanisms of initiation and cell of origin. We also identified an epigenetic signal associated with tumor burden, which may have some relation to viral infections such as Epstein-Barr-Virus. Our cell-type deconvolution-based approach to developing a continuous metric for CLL epigenetic differentiation state can be applied to other tumors with multiple subtypes across differentiation stages.

## 1 Introduction

Chronic Lymphocytic Leukemia (CLL) is a mature B-cell malignancy and the most frequent leukemia in the United States, occurring primarily in older adults [1]. CLL is clinically separated into two subgroups based on the mutational status of the immunoglobulin heavy-chain variable region (*IGHV*)[2, 3]. Cases with ≤98% identity to germline are deemed *IGHV* -mutated (M-CLL), indicating that the B-cell progenitor of the neoplasm has undergone somatic hypermutation; cases with *>* 98% identity are called *IGHV* -unmutated (U-CLL) [4]. This distinction has been validated as clinically significant for predicting risk of progression [3, 5] and response to therapies, [6, 7], and is concordant with multiple methods of molecular classification[8–10].

The cells of origin represented by U-CLL and M-CLL have been debated for over two decades [11, 12]. For U-CLL, the DNA-methylation profile has been previously characterized as B-naive like [10, 13], however other methods of DNA-methylation analyses report larger B-memory-like epigenetic signatures [14]. The cause of these discrepant characterizations remain unresolved, as well as the specific features of U-CLL which correspond to each normal cell reference.

Despite two tumor subtypes defined by *IGHV* mutational status, there exists a continuum of molecular states in the complex B-cell differentiation process that may represent different cell types of origin for CLL [14]. Previous work has used epigenetic signals to classify CLL into three epigenetic states [10, 15], or to assign CLL cases to their most similar cell of origin based on multiple references[16]. A continuous metric of differentiation for CLL based on the normal differentiation program has yet to be epigenetically explored and validated.

A key advancement in the analysis of bulk epigenetic data has been cell type deconvolution, which takes advantage of the distinct methylation signatures of purified cell types at specific CpG loci [17]. Recently, peripheral blood deconvolution libraries have been expanded to resolve 12 immune cell types [18], including distinguishing between the B-naive and B-memory cells. Deconvolution evaluates proportions within a mixture using constrained projection and a reference matrix of purified cell types [17]. While this approach was developed to quantify mixtures of discrete cell types, it has yet to be utilized for predicting intermediate states in purified abnormal cell populations.

Here, we develop a continuous metric for epigenetic B-differentiation state of CLL (B-Index) using novel applications of DNA methylation-based cell type deconvolution. B-Index is able to discriminate accurately between U-CLL and M-CLL, provides additional epigenetic information within subtypes, and carries stronger epigenetic signal than *IGHV* mutational status. Our normal reference-based approach also reveals that while U-CLL is more B-naive-like than M-CLL, it is more epigenetically similar to B-memory cells overall. These B-memory commonalities are enriched for many transcription factor binding sites related to B-cell activation, providing evidence for a post-activation cell of origin model for U-CLL. We also demonstrate that CLL has distinct epigenetic correlates to tumor burden.

## 2 Results

### 2.1 Development of B-Index

We collected peripheral blood samples from 89 patients with Chronic Lymphocytic Leukemia during their routine laboratory workups (Table 1). The study population consisted of 56 *IGHV* mutated (63%), 24 *IGHV* unmutated (27%), 3 Polyclonal *IGHV* (3%) and 6 cases where *IGHV* status was unavailable (7%). Subjects were predominately male (67%), and average age was 71.3. Information on *IGHV* mutational percentage was available for 46 participants.

**Table 1:** Cohort characteristics.

| Characteristic | Overall n = 89 |  |
| --- | --- | --- |
|  | Value | (%) |
| **Age at draw (years)** | 71.3 ± 9.1 | |
| **Sex** |  |  |
| Male | 60 | (67) |
| Female | 29 | (33) |
| ***IGHV* mutation status** |  |  |
| Mutated | 56 | (63) |
| Unmutated | 24 | (27) |
| Polyclonal | 3 | (3) |
| Failed | 6 | (7) |
| **Clonal B-cell count at diagnosis ($\times 10^3/\mu\text{L}$)** | 37.4 ± 68.0 | |
| **Sample lymphocyte count ($\times 10^3/\mu\text{L}$)** | 40.1 ± 74.2 | |

Continuous variables: mean ± SD. Categorical variables: *n* and (%).

CLL tumor cells were isolated from peripheral blood using a magnetic bead negative selection kit. DNA was extracted from isolated cells, quantified, and measured on high-density methylation microarrays. Quality control was performed on the methylation data, with all samples passing checks (Supplementary Figures 1A-D). 12 cell type deconvolution was performed using the Flow-Sorted Extended reference library to assess fractions of B-naive, B-memory, and non-B-cell signals. The deconvolution results consisted of almost entirely B-memory (Mean 0.81, SD 0.19) and B-naive (Mean 0.17, SD 0.19) fractions, showing very little signal from non B-cell fractions, with the largest being monocytes (Mean 0.018, SD 0.037) (Figure 1A-B).

**Figure 1:**
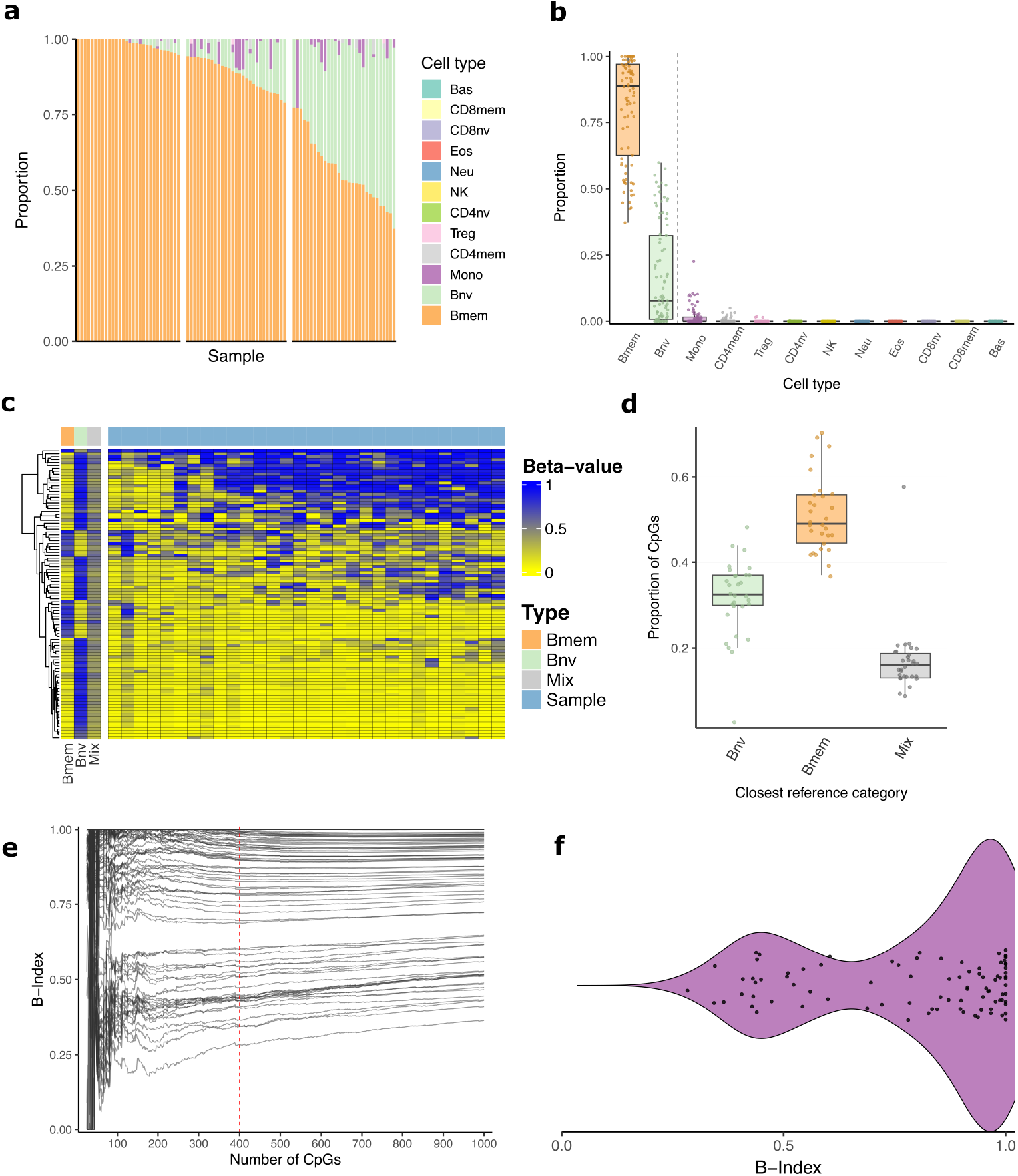
B-Index development. **a** Stacked bar plot representing results from 12 cell type epigenetic deconvolution, split into tertiles based on B-memory proportion. B-memory (Mean 0.81, SD 0.19) and B-naive (Mean 0.17, SD 0.19) show the largest proportions, with the largest non-B-cell population being Monocytes (Mean 0.018, SD 0.037). **b** Box plots of 12 cell-type deconvolution. Dashed vertical line separates B-cells from non-B-cells. **c** Heatmap of the top 100 most differentially methylated CpGs between B-memory and B-naive in the reference library. The samples with the most mixed predictions, taken from and in the same order as the right-most tertile of (a) (*n* = 30), are shown next to the B-memory and B-naive references, as well as an in-silico 60:40 mixture of B-memory and B-naive cells. **d** Box plots quantifying the number of CpGs per sample most similar to either i) pure B-naive reference ii) pure B-memory reference iii) a custom mixture based on each sample’s deconvolved proportions. Pure references explain the majority of CpGs (Mean 83%, SD 8.4%) in all samples except one. **e** Spaghetti plot showing the predicted B-Index of each sample as the number of CpGs included in prediction increases. CpGs were added in decreasing order of most differentially methylated between B-memory and B-naive within the reference. **f** Violin plot showing distribution of B-Index among all 89 samples.

As many of the samples had both B-naive and B-memory components, we sought to determine if the components represented either a single purified cell population that has individual epigenetic features of both B-Memory and B-naive cells, or a mixture of two distinct cell-type populations. To determine this, we tested the tertile of samples with the most mixed proportions (B-memory *<* 0.78, *n* = 30), and observed the top 100 most differentially methylated CpGs within the reference library between B-naive and B-memory, alongside an in-silico 60:40 mixture of B-memory and B-naive references. The samples were not similar to a mixture of two distinct cell populations (Figure 1C, Supplementary Figure 2). To quantify this, for each sample we created a custom mixed comparison group based on deconvolution-derived proportions and the reference library at the same 100 CpGs. For each sample in most mixed tertile, we compared Beta values of the sample to the custom mixture and the pure B-memory and B-naive references (Figure 1D). Results showed that for all the tested samples, more CpGs corresponded to a pure reference (Mean 83%, SD 8.4%) than to their custom mixture (Mean 17%, SD 8.4%), except for the sample assigned the largest monocyte proportion.

Having confirmed that the deconvolution results in this context constitute an intermediate state within a purified population, we sought to adapt the deconvolution framework to produce a continuous epigenetic surrogate of each sample’s B-cell differentiation state. To do this, non-B-cell references were removed from the 12 cell reference library, and the most differentially methylated CpGs between B-naive and B-memory were used for constrained projection. After performing constrained projection on a sample, the two outputted predictions of B-naive and B-memory were rescaled to sum to one. The B-Index is then defined as the predicted B-memory score after this rescaling. A score of 0 indicates high epigenetic similarity to B-naive, and a score of 1 indicates high similarity to B-memory. Samples can take on any continuous value across the 0-1 range. To determine how many CpGs to use for constrained projection, CpGs were added in order of most differentially methylated (Figure 1E). 400 was selected as a cutoff due to the stabilization of B-Index prediction. Among the full cohort, the distribution of B-Indexes was bimodal, with no sample having a B-Index lower than 0.28 (Figure 1F).

### 2.2 Validation and Epigenetic correlates of B-Index

B-Index values were significantly different between M-CLL (Mean 0.92, SD 0.094) and U-CLL (Mean 0.48, SD 0.096) subgroups by Wilcoxon rank-sum test (*p* = 2.6 × 10^−12^) (Figure 2A). To evaluate the utility of B-Index for inferring *IGHV* mutation status, we used it to classify samples into U-CLL and M-CLL. The area under the curve of the Receiver Operating Characteristic (ROC) was 0.995 (Figure 2B). An optimal 4 B-Index cutoff was found to be 0.60 by maximizing Youden’s J statistic. Using that threshold, B-Index was able to correctly classify samples into U-CLL and M-CLL with an accuracy of 98.8%. Samples without definitive IGHV mutational status information were excluded from classification analysis (Polyclonal *n* = 3, Failed *n* = 6). As U-CLL and M-CLL are categorical distinctions derived from tumor *IGHV* percent identity to germline, we compared B-Index to this underlying continuous variable. We found that *IGHV* mutational percentage and B-Index were correlated among the 46 (29 M-CLL and 17 U-CLL) samples where *IGHV* percentage was available (Supplementary Figure 3A), with an *R*^2^ of 0.65 (*p* = 1.1 × 10^−11^).

**Figure 2:**
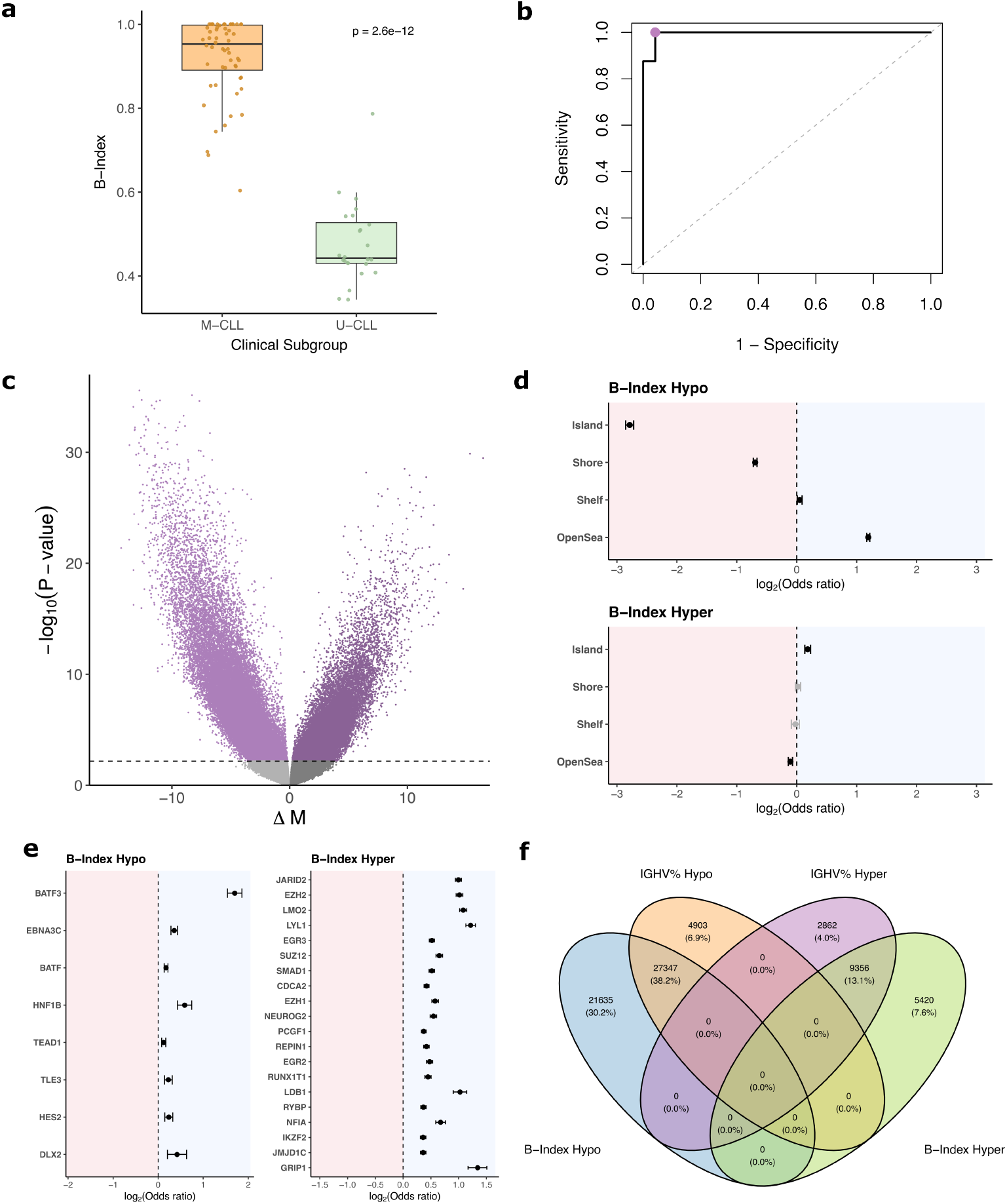
Validation and epigenetic correlates of B-Index. **a** Box plot of B-Index stratified into *IGHV* Mutated (M-CLL) and *IGHV* Unmutated (U-CLL). Mean B-Index of M-CLL was 0.92 (SD 0.094), mean B-Index for U-CLL was 0.48 (SD 0.096), the two distributions were significantly different by Wilcoxon test (*p* = 2.6 × 10^−12^). **b** Receiver Operating Characteristic (ROC) Curve for B-Index as a classifier into U-CLL and M-CLL. AUC was 0.995, and classification had 98.8% accuracy at Youden’s J optimal cutoff of 0.6. **c** Volcano plot depicting results from Epigenome Wide Association Study (EWAS) performed on B-Index. X-axis is delta M, or the change in M value per unit increase of the predictor (B-Index). The horizontal line marks the p-value corresponding to FDR adjusted *p <* 0.05. 83,129 CpGs were FDR significantly hypomethylated, and 30,710 hypermethylated. **d** Forest plot of island context enrichment among the hypo and hypermethylated CpGs with increasing B-Index, from (c). Odds ratio and 95% CI interval shown, in *log*_2_ space. Entries in black are significant (FDR *p <* 0.05). **e** Forest plot of transcription factor binding site enrichment among the hypo and hypermethylated CpGs with increasing B-Index, from (c). Odds ratio and 95% CI interval shown, in *log*_2_ space. Entries in black are significant (FDR *p <* 0.05). **f** Venn diagram of FDR significant CpGs from an EWASs conducted on *IGHV* mutational percent and on B-Index within the subset where *IGHV* mutational percent was available.

To determine the changes in the CLL methylome associated with B-Index, we used an epigenome wide association study (EWAS) with B-Index as our outcome variable. Using the full cohort (*n* = 89) and adjusting for the well documented epigenetic confounders of age and sex, increasing B-Index showed strong signals skewed toward hypomethylation, with 83,129 hypomethylated CpGs and 30,710 hypermethylated CpGs passing the false discovery rate (FDR) adjusted p-value threshold of 0.05 (Figure 2C) (Supplementary data 1). Among the genes related to the most significant hypomethylated CpGs were *LRRC31* (FDR *p* = 3.6 × 10^−30^), *ANO7* (FDR *p* = 1.1 × 10^−28^), and *OGG1* (FDR *p* = 3.6 × 10^−28^). Among the genes related to the most significant hypermethylated CpGs were *PRR33* (FDR *p* = 2.6 × 10^−26^) and *PPCDC* (FDR *p* = 3.7 × 10^−25^). Many of the most significant hypermethylated CpGs had multiple redundant gene hits, such as four CpGs related to *CRY1* (FDR p-values: 7.7 × 10^−23^, 2.0 × 10^−20^, 2.7 × 10^−20^, 2.3 × 10^−18^), two related to *LMAN2* (FDR p-values: 5.7 × 10^−26^, 2.4 × 10^−23^), and two related to *CLDN15* (FDR p-values: 6.6 × 10^−19^, 9.8 × 10^−19^).

To characterize the epigenetic signals correlated with B-Index at the hypomethylated and hypermethylated CpGs, we used enrichment analyses across multiple CpG attributes. We conducted a chromosomal enrichment analysis of the significantly (FDR p-value*<* 0.05) hypo and hypermethylated CpGs using Fisher’s exact test (Supplementary Figure 3B, Supplementary data 2-3). Among the hypomethylated CpGs, there was enrichment of chromosome 18 (OR= 1.20, FDR *p* = 2.7 × 10^−12^), 20 (OR= 1.15, FDR *p* = 3.9 × 10^−10^), and 21 (OR= 1.20, FDR *p* = 8.0 × 10^−9^) among others. Representation of chromosome 19 was significantly depleted (OR= 0.68, FDR *p* = 4.7 × 10^−89^), among others. The hypermethylated CpGs were enriched for chromosome 17 (OR= 1.27, FDR *p* = 6.2 × 10^−21^), among others. Chromosomes 2 (OR= 0.82, FDR *p* = 5.4 × 10^−17^) and 5 (OR= 0.80, FDR *p* = 4.7 × 10^−15^) were depleted, among others.

We also performed an enrichment analysis on CpG island relation, within the same sets of hyper/hypomethylated CpGs, adjusting for Illumina probe type using a Cochran-Mantel-Haenszel (CMH) test (Figure 2D, Supplementary data 4-5). Hypomethylated CpGs were depleted in CpG islands (OR= 0.14, FDR *p <* 10^−100^) and shore regions (OR= 0.62, FDR *p <* 10^−100^), and enriched in open-sea regions (OR= 2.3, FDR *p <* 10^−100^). Hypermethylated CpGs were enriched for representation of CpG islands (OR= 1.14, FDR *p* = 1.5 *×* 10^−13^), and depleted in open-sea (OR= 0.93, FDR *p* = 6.5 *×* 10^−9^). We further conducted an enrichment analysis on CpG genomic context, adjusting for island relation using a CMH Test (Supplementary Figure 3C, Supplementary data 6-7). Hypomethylated CpGs were enriched for intergenic (OR= 1.3, FDR *p <* 10^−100^) and intron regions (OR= 1.16, FDR *p* = 1.2 × 10^−82^), and depleted for promoter (OR= 0.81, FDR *p <* 10^−100^), 5’UTR (OR= 0.60, FDR *p* = 1.6 × 10^−49^), 3’UTR (OR= 0.68, FDR *p* = 1.0 × 10^−78^), and exon regions (OR= 0.84, FDR *p* = 8.3 × 10^−38^). Hypermethylated regions were enriched for 5’UTR (OR= 1.24, FDR *p* = 6.3 × 10^−11^) and intron regions (OR= 1.29, FDR *p* = 3.3 × 10^−93^), and depleted for promoter (OR= 0.82, FDR *p* = 1.7 × 10^−42^) and intergenic regions (OR= 0.69, FDR *p* = 4.5 × 10^−60^).

As transcription factor binding sites (TFBS) are known to change methylation states throughout many maturation pathways, we checked for enrichment of CpGs near TFBS within our CpGs that we found to be hypo/hypermethylated with increasing B-Index (Figure 2E, Supplementary data 8-9). Hypomethylated CpGs were enriched most significantly for binding sites of BATF3 (OR= 3.25, FDR *p* = 1.1 × 10^−76^). Hypermethylated CpGs were enriched in binding sites of JARID2 (OR= 1.99), EZH2 (OR= 2.02), LMO2 (OR= 2.12), and LYL1 (OR= 2.32) (all listed FDR*p <* 10^−100^), among others.

We also assessed enrichment among gene ontology (GO) terms. Hypomethylated CpGs showed enrichment for cell periphery (FDR *p* = 1.5 × 10^−43^), plasma membrane (FDR *p* = 4.0 × 10^−35^), and signaling receptor activity terms (FDR p= 2.0 × 10^−16^), among others (Supplementary Figure 3D, Supplementary data 10). An enrichment analysis of Kyoto Encyclopedia of Genes and Genomes (KEGG) terms showed significantly hypomethylated pathways in cornified envelope formation (FDR *p* = 1.72 × 10^−11^), among others (Supplementary Figure 3E, Supplementary data 11). For FDR significantly hypermethylated CpGs, KEGG analysis showed no significant results after FDR correction (Supplementary data 12). GO on the hypermethylated CpGs showed some FDR significant results, such as regulation of developmental processes (FDR *p* = 3.9 × 10^−3^) (Supplementary data 13).

To further assess the strength of the epigenetic correlates of B-Index in comparison to the underlying continuous clinical parameter, *IGHV* mutational status, we performed EWASs against B-Index and *IGHV* percentage germline identity on the subset of data where *IGHV* percent germline identity was available (*n* = 46, 29 M-CLL and 17 U-CLL cases), adjusting for age and sex (Supplementary Figure 4A-B, Supplementary data 14-15). The B-Index EWAS identified 14,776 significantly (FDR *p <* 0.05) hypermethylated CpGs, and 48,982 significantly (FDR *p <* 0.05) hypomethylated CpGs. *IGHV* mutational percent was significantly (FDR *p <* 0.05) associated with fewer CpGs than B-Index: 12,218 hypermethylated and 32,250 hypomethylated. The B-Index differentially methylated CpGs largely encompassed those found by the *IGHV* mutational percent EWAS (Figure 2F).

To determine whether continuous B-Index is able to identify additional epigenetic information within each subgroup, we stratified our cohort into U-CLL (*n* = 24) and M-CLL (*n* = 56) cases and performed an EWAS for B-Index on each. The EWAS on the U-CLL cases (Supplementary Figure 4C) found no FDR *p <* 0.05 significant hits. The EWAS on the M-CLL cases found 2,221 FDR *p <* 0.05 significantly hyper-methylated CpGs, and 16,459 significantly hypomethylated (Supplementary Figure 4D, Supplementary data 16). A Venn diagram of the B-Index associated CpGs from M-CLL and the B-Index associated CpGs from the full CLL cohort shows a partial overlap of the two signals (Supplementary Figure 4E).

### 2.3 U-CLL similarities to normal B-cell subsets

The U-CLL cases were found to have a mean B-Index of 0.48, indicating some commonalities with both B-naive and B-memory cells. We subsequently sought to further investigate the epigenetic similarities of U-CLL to B-naive and B-memory cells in terms of normal biology rather than CLL pathology. To serve as non-pathologic B-cell epigenome references, we used high density methylation microarray data (Illumina EPICv1 platform) consisting of 6 purified B-naive cell samples and 6 B-memory cell samples from healthy donors. Each cell population group was evenly split between male and female donors, with average ages of 38 (SD 10) for the B-naive donors and 32 (SD 12) for the B-memory donors.

We first determined the differentially methylated CpGs across the B-naive and B-memory cell populations, adjusting for the age and sex of the donors (Figure 3A, Supplementary data 17). After imposing a cutoff of FDR adjusted *p <* 0.05, and minimum change in beta value of 0.2, there were 108,978 significantly hypomethylated CpGs in B-memory compared to B-naive, and 11,156 significantly hypermethylated. Among the 120,134 differentially methylated CpGs, 97,709 were represented in the EPICv2 array and available in our CLL data set. Visualizing the top 1000 most significant differentially methylated CpGs between B-naive and B-memory cells in U-CLL cases showed strong similarity between U-CLL and B-memory cells (Figure 3B). In contrast, plotting the top 1000 most significantly differentially methylated CpGs from an EWAS comparing U-CLL to M-CLL (Supplementary Figure 5A, Supplementary data 18) shows the opposite pattern (Figure 3C), with the U-CLL cases appearing similar to B-naive cells. Visualizing all samples shows that U-CLL and M-CLL are both more memory-like than naive-like under the normal reference derived CpGs, and polarize to the references when using the CpGs differentially methylated between U-CLL and M-CLL (Supplementary Figure 6A-B).

**Figure 3:**
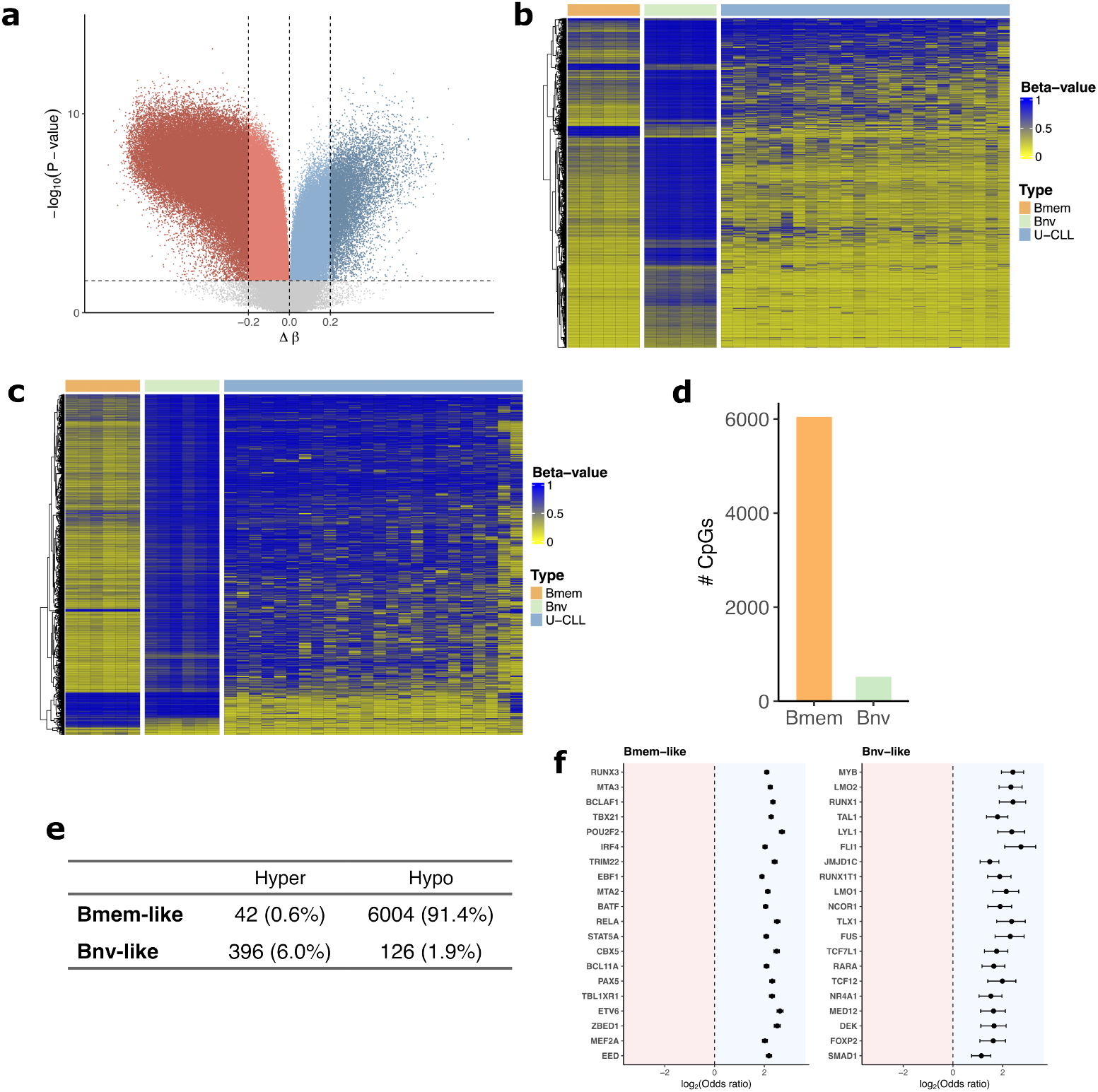
U-CLL similarities to B-memory and B-naive cells. **a** Volcano plot of EWAS between six purified B-memory references and six purified B-naive references. A cutoff of ± 0.2 Δ*β*, alongside the FDR *p <* 0.05 was used to determine differentially methylated CpGs between the two cell types. 108,978 CpGs were hypomethylated in B-memory with this criteria, and 11,156 were hypermethylated. **b** Heatmap of the top 1000 most significant CpGs from the B-memory v. B-naive EWAS available in our data. B-memory and B-naive references are shown alongside U-CLL samples ordered by increasing B-Index. **c** Heatmap similar to (b), except using top 1000 most significant CpGS from the U-CLL vs M-CLL EWAS. **d** Bar graph depicting results of two one-sided T (TOST) equivalence tests with acceptable equivalence bound at 0.1, assessing which CpGs in U-CLL are FDR *p <* 0.05 significantly equivalent to either B-memory or B-naive among the FDR significant and differentially methylated CpGs from (a). **e** Contingency table showing the methylation state of the TOST FDR *p <* 0.05 CpGs based on the normal B-cell to which they were equivalent in U-CLL, and their methylation state in U-CLL. **f** Forest plot of transcription factor binding site enrichment among CpGs in U-CLL similar (FDR *p <* 0.05) to B-memory and B-naive. Odds ratio and 95% CI interval shown. Entries in black are significant (FDR *p <* 0.05).

To quantify U-CLL’s epigenetic similarity to B-memory and B-naive cells, we performed equivalence tests between U-CLL and the normal B-cell groups on the 97,709 differentially methylated CpGs between B-memory and B-naive cells available in our data. We used the two-one-sided-T (TOST) test with an equivalence range of Beta +*/*−0.1. Among the beta values of CpGs tested, 6,046 were significantly (FDR p*<* 0.05) equivalent in U-CLL to B-memory cells, and 522 to B-naive cells (Figure 3D, Supplementary data 19). To verify that the B-memory like signature of U-CLL was not contingent on our CpG prefiltering, we computed the Manhattan distance of each U-CLL case to the mean Betas values among the B-memory and B-naive cases among all the 682,014 CpGs available across both datasets (Supplementary Figure 5B). Every U-CLL case had a higher Manhattan distance from B-naive than B-memory, indicating higher relative similarity with B-memory than B-naive.

To further characterize the B-naive and B-memory like signatures of U-CLL, we separated the FDR *p <* 0.05 CpGs from the TOST analysis into those that are in the hypermethylated or hypomethylated state in U-CLL (Figure 3E). Results showed that the U-CLL’s B-memory-like signature is hypomethylated, and its B-naive-like signature is mostly hypermethylated, with a Chi-squared test yielding a highly significant association (− log_10_(*p*) *>* 500).

In order to better discern in what particular ways U-CLL is B-memory and B-naive like, we conducted a series of enrichment analyses on both groups of the TOST equivalent CpGs. We used the TOST CpGs with FDR *p <* 0.05, and the 97,709 CpGs found to be different between B-memory and B-naive cells as the background to remove B-cell specific enrichment that is in common between B-memory and B-naive. Results for chromosomal enrichment were insignificant at all chromosomes for B-naive-like CpGs, and showed depletion at chromosome 5 in B-memory-like CpGs (OR= 0.80, FDR *p* = 5.1 × 10^−3^) (Supplementary data 20-21). CpG island relation enrichment was assessed using a CMH test, adjusting for Illumina probe type (Supplementary Figure 5C, Supplementary data 22-23). B-memory like CpGs were enriched for shore regions (OR= 1.25, FDR *p* = 3.5 × 10^−9^), and depleted in island (OR= 0.56, FDR *p* = 1.9 × 10^−8^) and open-sea (OR= 0.93, FDR *p* = 1.7 × 10^−2^) regions. B-naive like CpGs had no detectable enrichment signal. We then conducted enrichment analysis on genomic context, adjusting for island relation using a CMH Test (Supplementary Figure 5D, Supplementary data 24-25). B-memory like CpGs in U-CLL showed depletion in 3’UTR (OR= 0.71, FDR *p* = 6.8 × 10^−4^). B-naive like CpGs showed significant enrichment in 3’UTR (OR= 1.75, FDR *p* = 2.2 × 10^−2^) and exons (OR= 1.38, FDR *p* = 4.8 × 10^−2^), and depletion in intergenic regions (OR= 0.72, FDR *p* = 4.8 × 10^−2^). Enrichment of representation of GO and KEGG terms was also assessed among the B-memory and B-naive like CpGs, returning no *FDR <* 0.05 significant results (Supplementary data 26-29).

As transcription factor binding site (TFBS) methylation is known to change at many points along the normal B-naive to B-memory development, we assessed enrichment for TFBS among the B-memory and B-naive like CpGs in U-CLL (Figure 3F, Supplementary data 30-31). Among the many notable TFBS we found to be enriched among the CpGs similar to B-memory cells were POU2F2 (OR= 6.53), TBX21 (OR= 4.85), RELA (OR= 5.73), IRF4 (OR= 4.08), PAX5 (OR= 4.98), and BATF (OR= 4.15) (all listed FDR *p <* 10^−200^) among others. Outside of the top twenty shown in the figure were additional FDR-significant TFBS of interest, such as the NF-*κ*B family members: NFKB1 (OR= 3.54), NFKB2 (OR= 3.76), REL (OR= 5.90), and RELB (OR= 4.02), as well as BCL6 (OR= 7.25) and MEF2C (OR=6.45) (all listed FDR *p <* 10^−100^). Enriched binding sites among the CpGs similar to B-naive included LMO2 (OR= 5.03, FDR *p* = 3.3 *×* 10^−14^), RUNX1 (OR= 5.33, FDR *p* = 2.1 *×* 10^−11^), and FLI1 (OR= 6.66, FDR *p* = 2.0 *×* 10^−10^), among others.

### 2.4 Epigenetic correlates of disease burden

In addition to *IGHV* mutational data, for each study participant we also gathered two metrics of disease burden: the patient’s clonal B-cell count (× 10^3^ cells · *µ*L^−1^) at the time of their diagnosis (Mean 37 SD 68), and the total leukocyte count in sample, (10^3^ cells *µ*L^−1^) contemporary with their epigenetic data (Mean 40 SD 74) (Figure 4A). The two metrics, in *log*_2_ space, had *R*^2^ correlation of 0.53 (Supplementary Figure 7A). To assess the epigenetic correlates of each disease burden metric, EWASs were performed on total leukocyte count (Figure 4B, Supplementary data 32) and clonal B-cell count at diagnosis (Supplementary Figure 7B, Supplementary data 33) both in *log*_2_ space, adjusting for age and sex. 20,973 CpGs were found to be significantly (FDR p-value *<* 0.05) hypomethylated with increasing total leukocyte count, and 2,484 significantly hypermethylated. In line with the other burden metric, clonal B-cell count showed hypomethylation skewed epigenetic signature, with 16,475 significantly hypomethylated CpGs, and 997 hypermethylated. Based on total leukocytes count’s more significant epigenetic signal, it was used as the tumor burden metric of choice in all downstream analysis. To ensure this burden metric was independent from B-Index, a Pearson correlation test was performed across all samples on the *log*_2_ transformed burden metric, which showed no significant linear relationship (*R*^2^ = 0.004, *p* = 0.57) (Figure 4C). Adjusting for B-Index in the total leukocyte count EWAS also showed little change in significant CpGs (Supplementary Figure 7C, Supplementary data 34).

**Figure 4:**
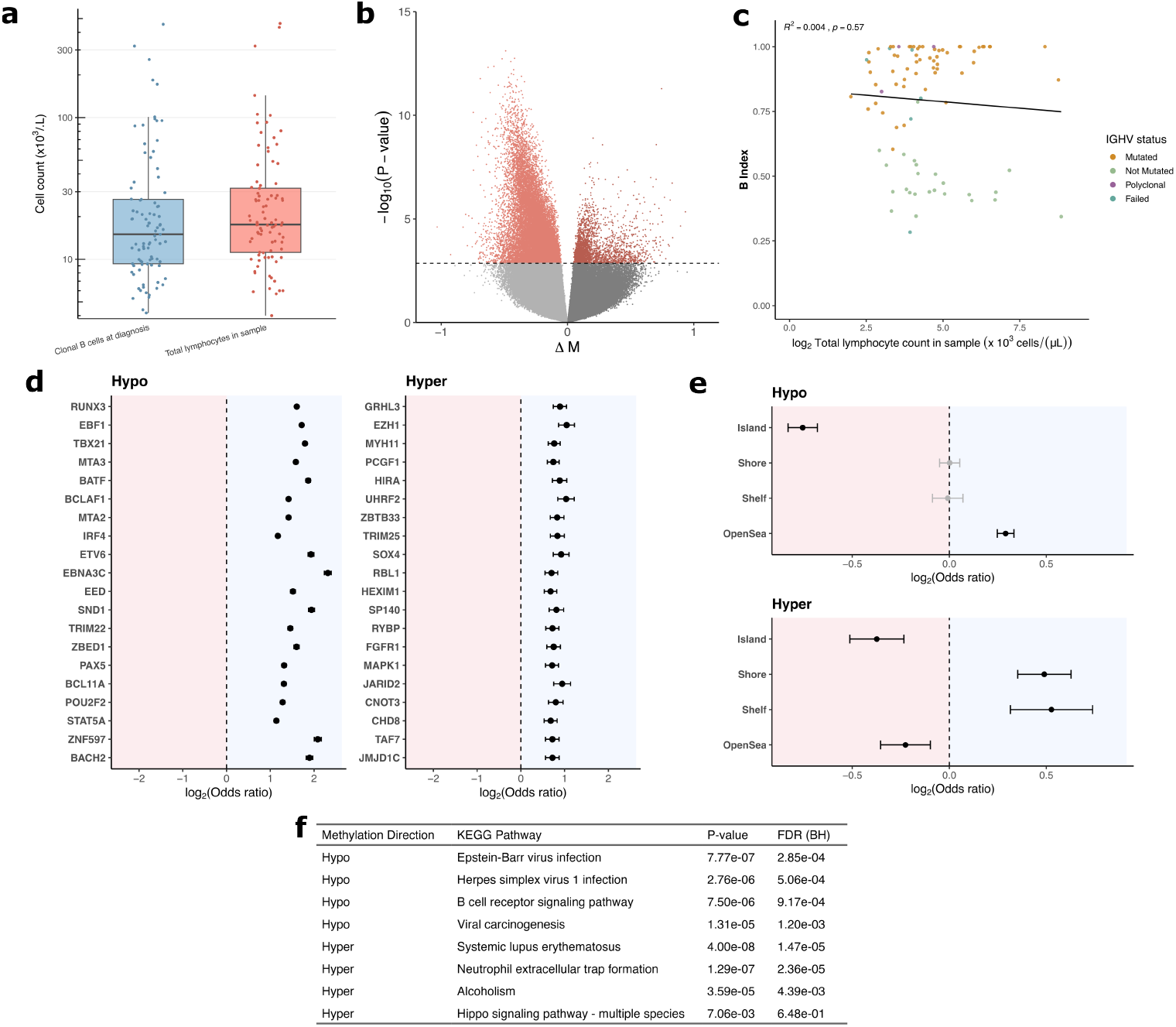
Epigenetic correlates of tumor burden. **a** Box plots depicting the two tumor burden metrics, in units of thousands of cells per microliter: clonal B-cell count at time of CLL diagnosis (Mean 37, SD 68), and total leukocyte count in sample (Mean 40 SD, 74). Y-axis is using log scaling. **b** EWAS on total leukocyte count, conducted in *log*_2_ space. 20,973 CpGs were FDR *p <* 0.05 significantly hypomethylated with increasing tumor burden, and 2,484 were FDR significantly hypermethylated. **c** Scatterplot of *log*_2_ total lymphocyte count and B-Index among the samples (*n* = 89, *R*^2^ = 0.004, *p* = 0.57). Colors of dots depict *IGHV* -mutational status. Line of best fit is shown. **d** Forest plot of transcription factor binding site enrichment among CpGs hypo and hypermethylated with increasing tumor burden. Odds ratio and 95% CI interval shown. Entries in black are significant (FDR *p <* 0.05) **e** Forest plot of CpG island context enrichment among CpGs hypo and hypermethylated with increasing tumor burden. Odds ratio and 95% CI interval shown. Entries in black are significant (FDR *p <* 0.05) **f** Top 4 hits from KEGG pathway analyses conducted separately on the hypermethylated and hypomethylated CpGs with increasing tumor burden.

To characterize the hypomethylation and hypermethylation signatures, we performed a series of enrichment analyses, beginning with chromosomal representation (Supplementary Figure 7D, Supplementary data 35-36). Hypermethylated CpGs were enriched for chromosome 13 (OR= 1.63, FDR *p* = 2.1 × 10^−4^), and depleted in chromosome 18 (OR= 0.61, FDR *p* = 2.7 × 10^−2^), among other significant deviations. Hypomethylated CpGs were enriched for chromosome 13 (OR= 1.29, FDR *p* = 3.6 × 10^−8^) and depleted in chromosome 7 (OR= 0.84, FDR *p* = 5.3 × 10^−7^), among others. We also assessed enrichment of CpG island context, adjusting for Illumina probe types using a CMH test (Figure 4D, Supplementary data 37-38). Hypomethylated CpGs were enriched for open-sea regions (OR= 1.22, FDR *p* = 1.5 × 10^−39^) and were depleted for island regions (OR= 0.59, FDR *p* = 8.5 × 10^−88^). Hypermethylated CpGs were enriched in shore (OR= 1.40, FDR *p* = 9.4 × 10^−12^) and shelf (OR= 1.44, FDR *p* = 1.3 × 10^−6^) regions, and depleted in open-sea (OR= 0.86, FDR *p* = 3.5 × 10^−4^) and island (OR= 0.77, FDR *p* = 8.7 × 10^−8^) regions. We then analyzed enrichment of genomic contexts using CMH tests adjusting for CpGs relationship to island (Supplementary Figure 7E, Supplementary data 39-40). No significant relationship was detected in the hypermethylated CpGs. The hypomethylated CpGs were enriched for intron regions (OR= 1.16, FDR *p* = 4.6 × 10^−24^), and depleted for intergenic (OR= 0.78, FDR *p* = 8.5 × 10^−23^), among others.

We then assessed enrichment of transcription factor binding sites (Figure 4E, Supplementary data 41-42). Hypomethylated CpGs were enriched for many sites of interest, including RUNX3 (OR= 3.04) EBF1 (OR= 3.29), BATF (OR= 3.65), EBNA3C (OR= 4.98) (all listed FDR *p <* 10^−300^), among others. Notably, EBNA3 (OR= 7.70, FDR *p <* 10^−100^), and EBNA2 (OR= 4.10, FDR *p* = 1.3 × 10^−67^), were also enriched. Hypermethylated regions were enriched in binding sites for EZH1 (OR= 2.06, FDR *p* = 1.9 × 10^−23^), UHRF2 (OR= 2.05, FDR *p* = 2.3 × 10^−21^), SOX4 (OR= 1.89, FDR *p* = 3.1 × 10^−19^), among others.

We further performed enrichment analyses on gene ontology (GO) terms and KEGG pathways (Supplementary Data 43-46). Among the significantly hypomethylated with increasing tumor burden, GO enrichment analysis revealed FDR significant hits for cytosol (FDR *p* = 4.1*×*10^−9^) and lymphocyte activation (FDR *p* = 5.2*×*10^−9^), among others (Supplementary Figure 7F). KEGG pathway analysis on hypomethylated CpGs revealed enrichment for the B cell receptor signaling pathway (FDR *p* = 9.2 × 10^−4^), Epstein-Barr virus infection (FDR *p* = 2.9 × 10^−4^), herpes simplex virus 1 infection (FDR *p* = 5.1 × 10^−4^) and viral carcinogenesis (FDR *p* = 1.2 × 10^−3^), among others (Figure 4F). Hypermethylated CpGs showed KEGG pathway enrichment of neutrophil extracellular trap formation (FDR *p* = 2.4 × 10^−5^) among others, and GO hits in structural constituent of chromatin (FDR *p* = 1.8 × 10^−4^), among others.

## 3 Discussion

We developed a continuous methylation-based metric for the B-differentiation state of CLL (B-Index) using a novel application of epigenetic cell type deconvolution. B-Index was able to discriminate between U-CLL and M-CLL with high accuracy. We also show that B-Index carries additional epigenetic correlates within the M-CLL sub-group, demonstrating the informative value that a continuous metric can provide, as noted previously [14]. B-Index carried a stronger epigenetic signal than the continuous metric underlying the clinical subgroup designation, *IGHV* mutational status. Higher B-Index was correlated mostly with hypomethylation, largely preferential to open-sea regions, and rarely occurring at CpG islands. Broad hypomethylation has been previously noted as characteristic of both B-cell differentiation, and CLL pathology [13, 19].

Hypomethylated CpGs associated with B-Index was also enriched near some TF binding sites, with the most significant being BATF3. BATF3 has been previously implicated in Anaplastic Large Cell Lymphoma as being overexpressed and essential to tumor growth [20]. Hypomethylation at BATF3 binding sites may increase chromatin accessibility or transcription factor binding ability [21]. Hypermethylation with B-Index occurred at binding sites near EZH2, which is the catalytic subunit in the Polychrome Repressor Complex 2 (PRC2) [22], and JARID2 which is known to recruit the PRC2 [23]. PRC2 activity has been shown to be up-regulated in other hematological malignancies, such as B-cell lymphomas [24]. EZH2 has also been implicated in lymphopoieses and immunoglobulin heavy chain rearrangement [25]. Hypermethylation also occurred at sites near LMO2 and LYL1 binding, both of which are known to play roles in lymphocyte differentiation [26, 27], and have been shown to have interactions essential to key features of Acute Lymphoblastic Leukemia (ALL) in mice [28].

In previous literature, U-CLL has often been described as epigenetically B-naive like [10, 13, 29], however, other previous studies have found the opposite characterization [14]. In this study, we are able to replicate both findings, dependent on the approach to data analysis. We find that when a common prefiltering of CpGs that are most differentially methylated between U-CLL and M-CLL is applied, U-CLL appears very B-naive-like. However, we also find that when prefiltering is performed in terms of normal tissue biology by selecting the CpGs most differentially methylated between B-naive and B-memory cells, U-CLL is strongly B-memory-like, with ∼ 12 fold more CpGs similar to B-memory cells than B-naive cells. We also find that when no prefiltering is applied, every one of our U-CLL cases remains more epigenetically similar to B-memory than B-naive. The origin of the different findings is that prefiltering CpGs to those most different or variable between or within a group of M-CLLs and U-CLLs removes their common epigenetic signal, which is B-memory-like.

We then characterized the specific CpGs where U-CLL is similar to B-memory and B-naive cells. We found that the B-memory-like CpGs in U-CLL are largely hypomethylated, and the B-naive-like are largely hypermethylated. This could be indicative of a majority, but incomplete transition to the B-memory epigenetic program, which mostly consists of hypomethylation [19]. In our transcription factor binding site analysis of B-memory similar CpGs in U-CLL, we found enrichment for many binding sites of TFs related to the germinal center response pathway [30], such as BCL6, MEF2C, IRF4, POU2F2 (OCT2), and NF-*κ*B family transcription factors (NFKB1, NFKB2, REL, RELB). We also demonstrated that these B-memory-like CpGs were overwhelmingly hypomethylated. Many of the B-memory-like binding sites identified to be B-memory-like in U-CLL in this study were shown previously to become hypomethylated between the states of naive and germinal center states of B-cells, and largely retain that hypomethylation through their development into B-memory cells (IRF4, MEF2C, MTA3, BATF, PAX5) (See TF enrichment on their M4 CpG module) [19].

Our analyses provide evidence for further defining cell type of origin and initiation mechanism for U-CLL. As we have demonstrated that U-CLL has many aspects of a B-memory-like signature, models of cell of origin should account for this in context with other known facts about U-CLL, such as its low levels of somatic hypermutation, and reconcile these through a common mechanism. Two possibilities include that the cell of origin of U-CLL has a largely B-memory-like methylation profile when it begins its transformation to a neoplasm, or U-CLL gains this methylation profile after becoming neoplastic. As we demonstrated in this study, a continuum across B-differentiation states can provide additional epigenetic information outside of the usual classification, which could indicate multiple underlying substates of origin among the already-used subtypes of CLL.

We also investigated epigenetic associations with tumor burden in CLL. In our samples, tumor burden was independent of B-Index and *IGHV* mutational status, and had an associated methylation signature. Higher tumor burden was correlated to ∼ 10 fold more hypomethylation than hypermethylation. Hypomethylation was enriched in open-sea regions and depleted in islands. The hypomethylated epigenetic signal was enriched for a number of transcription factor binding sites, including MTA2 and MTA3 which have been shown to have relationships to metastasis and tumor growth in multiple cancers [31–33]. Multiple Epstein-Barr-Virus (EBV) transcription factor binding sites were enriched among the hypomethylated CpGs as well, including EBNA3C, EBNA3, and EBNA2. KEGG terms were also enriched for CpGs related to EBV infection, as well as Herpes simplex virus 1 infection, and viral carcinogenesis. EBV has been shown to have an impact on time to treatment and overall survival in CLL [34], though the full relationship between EBV and CLL initiation and progression remains unclear [35]. Our data may provide motivation to analyze associations between the prescence of EBV in CLL cells and tumor growth and burden metrics for other researchers. As for the burden-correlated signals as a whole, with our study design we are unable to parse whether the epigenetic signal is variable over time and acquired as the CLL cell population grows, or precedes and predicts CLL clonal population growth.

Alongside our findings specific to CLL, we demonstrated a novel usage of DNA methylation-based cell-type deconvolution methods, applying constrained projection of intermediate cell states onto the purified cell population references. This approach may be useful in developing continuous metrics for other malignancies which can originate at multiple points along normal differentiation pathways, though this approach is not necessarily limited strictly to references on the same pathway. A reasonable skeleton for determining if this method is useful in another pathology would be to perform initial multi-cell-type deconvolution on a purified malignant population of interest. If the results yield a small number of dominating proportions, those can be further examined to develop a continuous index, as in this study.

## 4 Methods

### 4.1 Study Population, Sample Collection, and Clinical information

The 89 participants diagnosed with CLL were identified by clinicians during routine visits monitoring their disease. Participants consented to participate in the study, with IRB protocol number 020001881 approved by the Dartmouth-Hitchcock Medical Center committee for the protection of human subjects. Informed consent was obtained from all participants. Peripheral blood was collected from patients within the period of March 2023 to March 2024. Clinical information, such as *IGHV* mutational status and clonal B-cell count at diagnosis, was collected by chart review of participants. Total lymphocyte count in sample was determined by the CBC corresponding to the sample draw date.

### 4.2 Tumor Isolation and Methylation Measurement

After peripheral blood was collected from study participants, CLL cells were purified using immunomagnetic negative selection, with the EasySep(TM) Direct Human B-CLL Cell Isolation kit (Stemcell, Cat #19664), as per the kit protocol. The kit removes cells expressing CD2, CD3, CD14, CD16, CD56, CD61, CD66b. DNA was then extracted from the resulting purified cells, using EZ1 Advanced XL DNA extraction platform (Qiagen, cat #9001875), using the blood card. DNA underwent Bisulfite conversion using Zymo EZ DNA Methylation Kit, and was measured using Illumina EPICv2 array based methylation platform, according to manufacturer protocol.

### 4.3 Methylation Preprocessing and Quality Control

Methylation data, was preprocessed using the minfi (version 1.52.1)[36], Sesame (version 1.24.0)[37], and ENmix (version 1.42.2)[38] packages in R. Sex was predicted using the median raw probe intensities on the X and Y chromosomes, using the minfi getSex function. Predicted sex was matched to labeled sex to identify any potentially mislabeled epigenetic data. Low quality probes, meaning probes that had detection P-values *>* 10^−6^ or less than three beads available in more than 5% of samples, were identified using ENmix’s QCinfo function. All samples had less than 5% of their probes marked as low quality and were within three standard deviations of the mean bisulfite conversion intensity, and were retained. Beta values were generated through the Normal-Exponential-Out-of-Band (NOOB) preprocessing method using minfi, in order to account for dye-bias. Probes annotated to the X and Y chromosome were filtered out, based on Illumina’s 20a1.hg38 annotation file. SNP and cross-hybridizing probes, were filtered out using the EPICv2 mask general file from zwdzwd.github.io/InfiniumAnnotation. Probes not corresponding to CpGs, or not included in the illumina annotation file were also filtered. Technical replicate probe Beta values were collapsed to their averages using the betasCollapseToPfx function from sesame R package. Beta distribution graphs were generated by Gaussian kernel density estimation, using Sheather-Jones bandwidth selector, and 512 bins. After probe filtering and collapsing replicates 843,782 CpGs remained in the dataset. M values, which were used for epigenome wide assocation studies (EWAS) in order to better fulfill homoscedasticity assumptions in linear modeling, were calculated using the following equation:

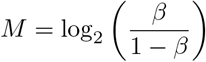

### 4.4 DNA Methylation Based Deconvolution

DNA-methylation based cell type deconvolution was conducted using a 12-cell reference library [18]. The library contains reference beta values for 12 Cell types (Basophils, B-memory, B-naive, CD4-memory, CD4-naive, CD8-memory, CD8-naive, Eosinophils, Monocytes, Neutrophils, Natural Killers, and T-regulators), at 1,200 CpGs selected through the Identifying Optimal Libraries (IDOL) algorithm. Library was subsetted to CpGs that overlapped with those available in our preprocessed data. Constrained projection, as implemented in the function projectCellType CP in the FlowSorted.BloodExtended.EPIC package (version 0.99.0) [39], was applied on the reference library matrix and the CLL methylation data to produce predicted proportions. Predicted proportions were normalized to sum to one per sample.

### 4.5 Weighted Reference Mixture Modeling

To determine if the deconvolution results corresponded better to a mixture of two distinct cell populations, or a single population with individual CpG methylation states of two references (B-memory and B-naive), we created custom references for each mixed sample at each CpG. The samples were divided into tertiles based on B-memory proportion, and the tertile with the smallest B-memory proportions (*n* = 30, B-memory *<* 0.78) was tested. The top 100 most differentially methylated CpGs between the B-naive and B-memory reference available in our data were used. For each mixed sample and CpG, we estimated the expected beta value as the weighted average of reference betas, using the sample’s predicted cell-type proportions as weights. This yields the beta value expected if the sample is a true mixture rather than a single population with distinct epigenetic features of B-memory and B-naive cells. For a single sample, the following equation was used to generate the sample’s mixed reference.

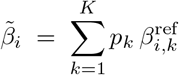

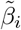 represents the predicted beta value at CpG *i* for sample given the mixture of cell types, *K* is the number of reference cell types, *p*_*k*_ is the predicted proportion for cell type *k*. 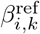 is the reference beta value at CpG *i* for cell type *k, i* is the index for each of the 100 CpG sites, and is the index for *k* reference cell types.

The actual Beta value for each sample and CpG was then compared to the beta value of the predicted mixture, the purified B-memory reference, and the purified B-naive reference to see which it was most similar to by absolute difference. The CpG was then tallied to the most similar comparison group.

To visualize the mixture reference a 60:40 weighted average of the reference B-memory and B-naive CpGs were used, based on the mean proportions of the predicted mixed samples.

### 4.6 B-Index

To develop B-Index, the prediction score for CLL where 0 corresponds to epigenetically B-naive-like, and 1 corresponds to epigenetically B-memory like, techniques based on DNA methylation cell type deconvolution were used. The 12 cell reference library described above was subsetted to just the B-memory and B-naive references, with the top most differentially methylated CpGs between the two references available in our data included. Constrained projection was then performed using the projectCell-Type CP function from the FlowSorted.BloodExtended.EPIC package (version 0.99.0) [39]. The two resulting proportions were restricted to a minimum of 0. The results were then normalized to sum to 1. The B-Index was then defined to be the B-memory proportion after the previously described operations. The equation for B-Index for a sample given predicted B-memory proportion 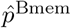 and predicted B-naive proportion 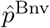 is shown below:

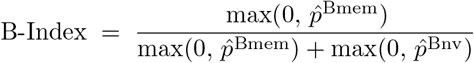

To determine how many of the top most differentially methylated CpGs between the B-naive and B-memory references to include, different amounts from 10 to the total amount available in our data from the reference library were used to calculate B-Index for our samples. After visualization of the resulting B-Indexes on a spaghetti plot, 400 was selected due to the stabilization of B-Index variation per sample over increasing amounts of included CpGs.

### 4.7 Epigenome Wide Association Studies

Multiple Epigenome Wide Association Studies (EWASs) were conducted. EWASs are linear regression models of CpG Beta-values or M-values with respect to input parameters. For every EWAS in this study, M-values, which are defined as 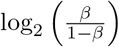 as described above, were used as the modeled variable in order to better fulfill the assumption of homoscedasticity, which B-value distributions tend to violate due to their bounding. For every EWAS, known confounders of epigenetic information, age and sex, were adjusted for by inclusion in the model matrix. Age was centered around the mean value in model matrix. The EWAS outputs linear models of the M-values, where Beta-Coeffecients of variables of interest and adjusting variables are estimated using linear regression. The resulting Beta-Coefficient of the variable of interest (such as B-Index) is labeled as Δ*M*, which can be interpreted as the estimated change in M-value per unit increase of variable of interest. Positive Δ*M* corresponds to Hypermethylation at given CpG with respect to variable of interest, and negative Δ*M* corresponds to hypomethylation.

For one EWAS, a transformation from Δ*M* to Δ*β* was conducted in order to apply a Δ*β* cutoff. To do this, the following equation was used (as described previously [40]), requiring the Δ*M* (Beta coefficient of variable of interest, when M-Values used as output), and the intercept of the regression *β*_0_:

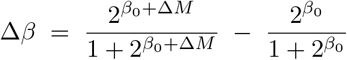

EWAS were implemented using the limma package (version 3.62.2)[41], using the lmFit function to perform regressions and the eBayes function to perform empirical Bayes moderation of standard errors. Adjusted p-values were calculated using Benjamini-Hochberg false discovery rate method, with all inputted CpGs being used in the corrections.

### 4.8 Enrichment Analyses

A number of enrichment studies, which assess whether a given list of CpGs have an overrepresentation of features of interest when compared to a background, were performed in this study. Enrichment of Gene Onotology (GO) terms was performed to assess if CpGs of interest were enriched for functional gene groups. Kyoto Encyclopedia of Genes and Genomes (KEGG) analysis was performed to assess enrichment for biological pathways. GO and KEGG analysis were performed using the gometh function in the MissMethyl package (version 1.40.3)[42], using the 20a1.hg38 Illumina Epic V2 annotation (version 1.0.0)[43]. MissMethyl accounts for bias in both probes per gene on the EPICv2 array, and CpG annotation to multiple genes.

For CpG island context and chromosomal enrichment, the 20a1.hg38 Illumina Epic V2 annotation file was used to map CpGs. Two-sided Fisher’s exact test was used for chromosomal enrichment. For genomic context, annotation was constructed using AnnotationHub (version 3.14.0)[44], and Ensembl version 113, genomic ranges of exons, introns, 5’UTR, 3’UTR, and promoters were accessed and mapped to probe coordinates using sesameData getManifestGRanges. Promoter range was defined as up to 1,500 bases upstream, and up to 200 bases downstream. Probes with multiple mappings were collapsed to a single genomic context using the following order of precedence: Promoter *>* 5’UTR *>* 3’UTR *>* Exon *>* Intron. Probes not mapping to any genomic ranges were labeled intergenic. As CpG islands associate with genomic context, a Cochran-Mantel-Haenszel (CMH) test, stratifying across CpG island context, was used. Likewise, island context enrichment was tested using a CMH test, stratifying across Illumina probe type (Type I/Type II).

For transcription factor binding site analysis, the KnowYourCG (KYCG) package (version 1.2.5)[45] EPICv2 consensus transcription factor binding site (TBFS) knowledge base was used. KYCG consensus TBFS annotations are constructed by integrating ChIP-seq peaks from Cistrome and ReMap (derived from ENCODE), and ranking CpGs by overlap frequency among multiple datasets to capture consensus binding features. We collapsed this annotation to probe prefixes to match with our probe annotation, and removed resulting duplicates. Enrichment tests were then conducted with KnowYourCG’s testEnrichment function, set to alternative=“greater”, which uses a one-sided Fisher’s exact test. Confidence intervals were added to functions output using the same statistical method.

For all enrichment tests, cutoff of BH adjusted FDR P value *<* 0.05 was used. For all EWAS derived enrichment analyses, all EWAS inputted CpGs were used as the total list, and CpGs were split into hypermethylated and hypomethylated for separate analyses. For the enrichment tests on CpGs determined to be similar to either B-naive or B-memory in U-CLL by TOST, all CpGs inputted into the TOST were used as the total list, in order to control for enrichment that B cell specific CpGs have in tested features, and enrichment analyses were performed separately on B-memory-like and B-naive-like CpGs. FDR p-values were calculated based on number of tests conducted per analysis. Odds ratios with 95% confidence intervals were reported, as well as FDR BH adjusted P-values. Forest plots are in *log*_2_ space.

### 4.9 Normal B-Cell References

The twelve normal B-cell references (6 B-memory, 6 B-naive) were acquired via Gene Expression Omnibus (GEO) accession GSE174666. As described previously [46], the B-memory and B-naive cells were isolated via magnetic activated cell sorting (MACS) techniques. Leukocytes from adult whole blood were separated into CD27+ and CD27−, and then CD19+ and CD19−. B-memory cell populations (CD19+ and CD27+), and B-naive cell populations (CD19+ and CD27−), were reported to have purities between 90-98%. Extracted DNA was measured on Illumina EPICv1 microarray, and preprocessing was performed as described in the original study[46]. The processed Beta matrix was retrieved from the GEO accession, along with phenotypic data. The beta values were converted to M-values, as described above.

### 4.10 Equivalence Testing

To test equivalence between methylation states in U-CLL and the B-memory and B-naive references, we used the two one-sided T-Tests (TOST) method, implemented in the TOSTER package (version 0.8.4)[47]. Among the results from the EWAS on the normal B-references, CpGs with FDR *p >* 0.05 and |Δ*β*| *>*0.2 that were also available in our data were used for the analysis. At each of these CpGs, the U-CLL samples were tested for TOST equivalence to the B-naive and B-memory using an epsilon of 0.1, meaning a ± 0.1 difference in Beta value will be tolerated for equivalence. As standard with TOST analysis, the larger of the two p-values from the one sided tests was taken as the p-value for the TOST. Benjamini-Hochberg FDR corrected p-values were calculated based on total CpGs inputted to the TOST.

### 4.11 Additional Statistical Analysis

Analyses were performed on R (version 4.4.1)[48]. Wilcoxon rank sum test was performed using the wilcox.test function, without continuity correction. Receiver operating curves (ROCs), and Youden’s J statistics were generated using the PRROC package (version 1.4)[49].

## 5 Data availability

Epigenetic and phenotypic data from the 89 Samples using this study, including raw .IDAT files, the preprocessed beta matrix, B-Index, *IGHV* mutational status, *IGHV* mutational percentage, absolute clonal B-cell count at diagnosis, total lymphocyte count in sample, age, and sex are deposited on Gene Expression Omnibus (GEO), available upon publication. Ages at 90 or above are bucketed and marked on GEO. Epigenetic and phenotypic data for the normal B-naive and B-memory references are available at Gene Expression Omnibus at accession GSE174666 [46].

## 6 Code availability

All code used in analyses, figure generation, and all other code related to this manuscript are deposited on Zenodo at 10.5281/zenodo.17118029.

## 7 Author contributions

J.H conceived of study question, designed and carried out analysis plan, and wrote the manuscript with input from all authors. B.C. assisted in refinement of analysis plan. B.C. and P.K. supervised the project, designed initial study purpose, and acquired funding. L.W. conducted sample collection, sample processing, assisted in grant writing, and acquired clinical information. D.S. assisted in acquiring clinical information. S.S. assisted in sample processing. F.K.IV conducted methylation data collection. F.K. and P.P. contributed to data processing. All authors provided feedback throughout the study and approved the final version.

## 8 Acknowledgments

The authors acknowledge the support of the Clinical Genomics and Advanced Technology Section in the Department of Pathology and Laboratory Medicine of the Dartmouth Hitchcock Health System, and the Pathology Shared Resource (RRID: SCR 023479) at the Dartmouth Cancer Center with NCI Cancer Center Support Grant 5P30 CA023108-41. Methylation microarray measurement was carried out in the Genomics and Molecular Biology Shared Resource (RRID:SCR 021293) at Dart-mouth which is supported by NCI Cancer Center Support Grant 5P30CA023108 and NIH S10 (1S10OD030242) awards. The authors also acknowledge pilot funding from the Dartmouth Cancer Center.

## References

[1] Hallek, M. Chronic Lymphocytic Leukemia: 2025 Update on the Epidemiology, Pathogenesis, Diagnosis, and Therapy. American Journal of Hematology 100, 450–480 (2025). URL https://onlinelibrary.wiley.com/doi/abs/10.1002/ajh.27546. eprint: https://onlinelibrary.wiley.com/doi/pdf/10.1002/ajh.27546.

[2] Damle, R. N. et al. Ig V Gene Mutation Status and CD38 Expression As Novel Prognostic Indicators in Chronic Lymphocytic Leukemia: Presented in part at the 40th Annual Meeting of The American Society of Hematology, held in Miami Beach, FL, December 4-8, 1998. Blood 94, 1840–1847 (1999).

[3] Hamblin, T. J., Davis, Z., Gardiner, A., Oscier, D. G. & Stevenson, F. K. Unmutated Ig VH Genes Are Associated With a More Aggressive Form of Chronic Lymphocytic Leukemia. Blood 94, 1848–1854 (1999).

[4] Ghia, P. et al. ERIC recommendations on IGHV gene mutational status analysis in chronic lymphocytic leukemia. Leukemia 21, 1–3 (2007).

[5] Galieni, P. et al. Unmutated IGHV at diagnosis in patients with early stage CLL independently predicts for shorter follow-up time to first treatment (TTFT). Leukemia Research 143, 107541 (2024).

[6] Rotbain, E. C. et al. IGHV mutational status and outcome for patients with chronic lymphocytic leukemia upon treatment: A Danish nationwide populationbased study. Haematologica 105, 1621–1629 (2020).

[7] Visentin, A. et al. Prognostic and Predictive Effect of IGHV Mutational Status and Load in Chronic Lymphocytic Leukemia: Focus on FCR and BR Treatments. Clinical Lymphoma Myeloma and Leukemia 19, 678–685.e4 (2019).

[8] Knisbacher, B. A. et al. Molecular map of chronic lymphocytic leukemia and its impact on outcome. Nature genetics 54, 1664–1674 (2022).

[9] Li, F. J. et al. FCRL2 expression predicts IGHV mutation status and clinical progression in chronic lymphocytic leukemia. Blood 112, 179–187 (2008).

[10] Queirós, A. C. et al. A B-cell epigenetic signature defines three biologic subgroups of chronic lymphocytic leukemia with clinical impact. Leukemia 29, 598–605 (2015).

[11] Caligaris-Cappio, F. & Ghia, P. The nature and origin of the B-chronic lymphocytic leukemia cell: A tentative model. Hematology/Oncology Clinics 18, 849–862 (2004).

[12] Chiorazzi, N. & Ferrarini, M. Cellular origin(s) of chronic lymphocytic leukemia: Cautionary notes and additional considerations and possibilities. Blood 117, 1781–1791 (2011).

[13] Kulis, M. et al. Epigenomic analysis detects widespread gene-body DNA hypomethylation in chronic lymphocytic leukemia. Nature Genetics 44, 1236– 1242 (2012).

[14] Oakes, C. C. et al. DNA methylation dynamics during B cell maturation underlie a continuum of disease phenotypes in chronic lymphocytic leukemia. Nature Genetics 48, 253–264 (2016).

[15] Giacopelli, B. et al. Developmental subtypes assessed by DNA methylation-iPLEX forecast the natural history of chronic lymphocytic leukemia. Blood 134, 688–698 (2019).

[16] Wierzbinska, J. A. et al. Methylome-based cell-of-origin modeling (Methyl-COOM) identifies aberrant expression of immune regulatory molecules in CLL. Genome Medicine 12, 29 (2020).

[17] Houseman, E. A. et al. DNA methylation arrays as surrogate measures of cell mixture distribution. BMC Bioinformatics 13, 86 (2012).

[18] Salas, L. A. et al. Enhanced cell deconvolution of peripheral blood using DNA methylation for high-resolution immune profiling. Nature Communications 13, 761 (2022).

[19] Kulis, M. et al. Whole-genome fingerprint of the DNA methylome during human B cell differentiation. Nature Genetics 47, 746–756 (2015). URL https://www.nature.com/articles/ng.3291. Publisher: Nature Publishing Group.

[20] Schleussner, N. et al. The AP-1-BATF and -BATF3 module is essential for growth, survival and TH17/ILC3 skewing of anaplastic large cell lymphoma. Leukemia 32, 1994–2007 (2018). URL https://www.nature.com/articles/s41375-018-0045-9. Publisher: Nature Publishing Group.

[21] Kaluscha, S. et al. Evidence that direct inhibition of transcription factor binding is the prevailing mode of gene and repeat repression by DNA methylation. Nature Genetics 54, 1895–1906 (2022). URL https://www.nature.com/articles/s41588-022-01241-6. Publisher: Nature Publishing Group.

[22] Margueron, R. & Reinberg, D. The Polycomb complex PRC2 and its mark in life. Nature 469, 343–349 (2011). URL https://www.nature.com/articles/nature09784. Publisher: Nature Publishing Group.

[23] Li, G. et al. Jarid2 and PRC2, partners in regulating gene expression. Genes & Development 24, 368–380 (2010). URL http://genesdev.cshlp.org/content/24/4/368. Company: Cold Spring Harbor Laboratory Press Distributor: Cold Spring Harbor Laboratory Press Institution: Cold Spring Harbor Laboratory Press Label: Cold Spring Harbor Laboratory Press Publisher: Cold Spring Harbor Lab.

[24] McCabe, M. T. et al. Mutation of A677 in histone methyltransferase EZH2 in human B-cell lymphoma promotes hypertrimethylation of histone H3 on lysine 27 (H3K27). Proceedings of the National Academy of Sciences 109, 2989– 2994 (2012). URL https://www.pnas.org/doi/full/10.1073/pnas.1116418109. Publisher: Proceedings of the National Academy of Sciences.

[25] Su, I.-h. et al. Ezh2 controls B cell development through histone H3 methylation and Igh rearrangement. Nature Immunology 4, 124–131 (2003). URL https://www.nature.com/articles/ni876. Publisher: Nature Publishing Group.

[26] Hirano, K.-i. et al. LMO2 is essential to maintain the ability of progenitors to differentiate into T-cell lineage in mice. eLife 10, e68227 (2021). URL 10.7554/eLife.68227. Publisher: eLife Sciences Publications, Ltd.

[27] Capron, C. et al. The SCL relative LYL-1 is required for fetal and adult hematopoietic stem cell function and B-cell differentiation. Blood 107, 4678–4686 (2006). URL 10.1182/blood-2005-08-3145.

[28] McCormack, M. P. et al. Requirement for Lyl1 in a model of Lmo2-driven early T-cell precursor ALL. Blood 122, 2093–2103 (2013). URL 10.1182/blood-2012-09-458570.

[29] Turk, A., Čeh, E., Calin, G. A. & Kunej, T. Multiple omics levels of chronic lymphocytic leukemia. Cell Death Discovery 10, 293 (2024). URL https://www.nature.com/articles/s41420-024-02068-2. Publisher: Nature Publishing Group.

[30] Laidlaw, B. J. & Cyster, J. G. Transcriptional regulation of memory B cell differentiation. Nature Reviews. Immunology 21, 209–220 (2021). URL https://www.ncbi.nlm.nih.gov/pmc/articles/PMC7538181/.

[31] Zhou, C. et al. MTA2 promotes gastric cancer cells invasion and is transcriptionally regulated by Sp1. Molecular Cancer 12, 102 (2013). URL 10.1186/1476-4598-12-102.

[32] Lin, C.-L. et al. MTA2 silencing attenuates the metastatic potential of cervical cancer cells by inhibiting AP1-mediated MMP12 expression via the ASK1/MEK3/p38/YB1 axis. Cell Death & Disease 12, 451 (2021). URL https://www.nature.com/articles/s41419-021-03729-1. Publisher: Nature Publishing Group.

[33] Dong, H. et al. The metastasis-associated gene MTA3, a component of the Mi-2/NuRD transcriptional repression complex, predicts prognosis of gastroesophageal junction adenocarcinoma. PloS One 8, e62986 (2013).

[34] Liang, J.-H. et al. Prognostic impact of Epstein-Barr virus (EBV)-DNA copy number at diagnosis in chronic lymphocytic leukemia. Oncotarget 7, 2135–2142 (2015). URL https://www.ncbi.nlm.nih.gov/pmc/articles/PMC4811522/.

[35] Dolcetti, R. & Carbone, A. Epstein-Barr virus infection and chronic lymphocytic leukemia: a possible progression factor? Infectious Agents and Cancer 5, 22 (2010). 10.1186/1750-9378-5-22.

[36] Aryee, M. J. et al. Minfi: a flexible and comprehensive Bioconductor package for the analysis of Infinium DNA methylation microarrays. Bioinformatics (Oxford, England) 30, 1363–1369 (2014).

[37] Zhou, W., Triche, T. J., Jr, Laird, P. W. & Shen, H. SeSAMe: reducing artifactual detection of DNA methylation by Infinium BeadChips in genomic deletions. Nucleic Acids Research 46, e123 (2018). URL 10.1093/nar/gky691.

[38] Xu, Z., Niu, L. & Taylor, J. A. The ENmix DNA methylation analysis pipeline for Illumina BeadChip and comparisons with seven other preprocessing pipelines. Clinical Epigenetics 13, 216 (2021). URL 10.1186/s13148-021-01207-1.

[39] Salas, L. A. & Zhang, Z. FlowSorted.BloodExtended.EPIC: Illumina EPIC data on extended immunomagnetic sorted blood populations (2021). URL https://github.com/immunomethylomics/FlowSorted.BloodExtended.EPIC. R package version 0.99.0, commit 1a2e43ae956e8d886b672f1098d9f363c272549e.

[40] Xie, C. et al. Differential methylation values in differential methylation analysis. Bioinformatics 35, 1094–1097 (2019). URL https://www.ncbi.nlm.nih.gov/pmc/articles/PMC6449748/.

[41] Ritchie, M. E. et al. limma powers differential expression analyses for RNA-sequencing and microarray studies. Nucleic Acids Research 43, e47 (2015).

[42] Phipson, B., Maksimovic, J. & Oshlack, A. missMethyl: an R package for analyzing data from Illumina’s HumanMethylation450 platform. Bioinformatics 32, 286–288 (2016). URL 10.1093/bioinformatics/btv560.

[43] Gu, Z. IlluminaHumanMethylationEPICv2anno.20a1.hg38: Annotation for Illumina’s EPIC v2.0 methylation arrays (2024). URL https://github.com/jokergoo/IlluminaHumanMethylationEPICv2anno.20a1.hg38. R package version 1.0.0, commit 702e52acb8268353bd8c753e67a2096c565f9782.

[44] Morgan, M. & Shepherd, L. AnnotationHub: Client to access AnnotationHub resources (2024). URL https://bioconductor.org/packages/AnnotationHub. R package version 3.14.0.

[45] Wanding, Z. & David, G. knowYourCG: Functional analysis of DNA methylome datasets (2024). URL https://bioconductor.org/packages/knowYourCG. R package version 1.2.5.

[46] Zhang, Z. et al. Comparative analysis of the DNA methylation landscape in CD4, CD8, and B memory lineages. Clinical Epigenetics 14, 173 (2022).

[47] Lakens, D. Equivalence Tests: A Practical Primer for t Tests, Correlations, and Meta-Analyses. Social Psychological and Personality Science 8, 355–362 (2017). URL 10.1177/1948550617697177. Publisher: SAGE Publications Inc.

[48] R Core Team. R: A Language and Environment for Statistical Computing. R Foundation for Statistical Computing, Vienna, Austria (2024). URL https://www.R-project.org/.

[49] Keilwagen, J., Grosse, I. & Grau, J. Area under Precision-Recall Curves for Weighted and Unweighted Data. PLOS ONE 9, e92209 (2014). URL https://journals.plos.org/plosone/article?id=10.1371/journal.pone.0092209. Publisher: Public Library of Science.

