## Supplemental Figures for "Continuous DNA Methylation Deconvolution-Based Surrogate for B-Cell Differentiation State in CLL"

### Supplementary Figures

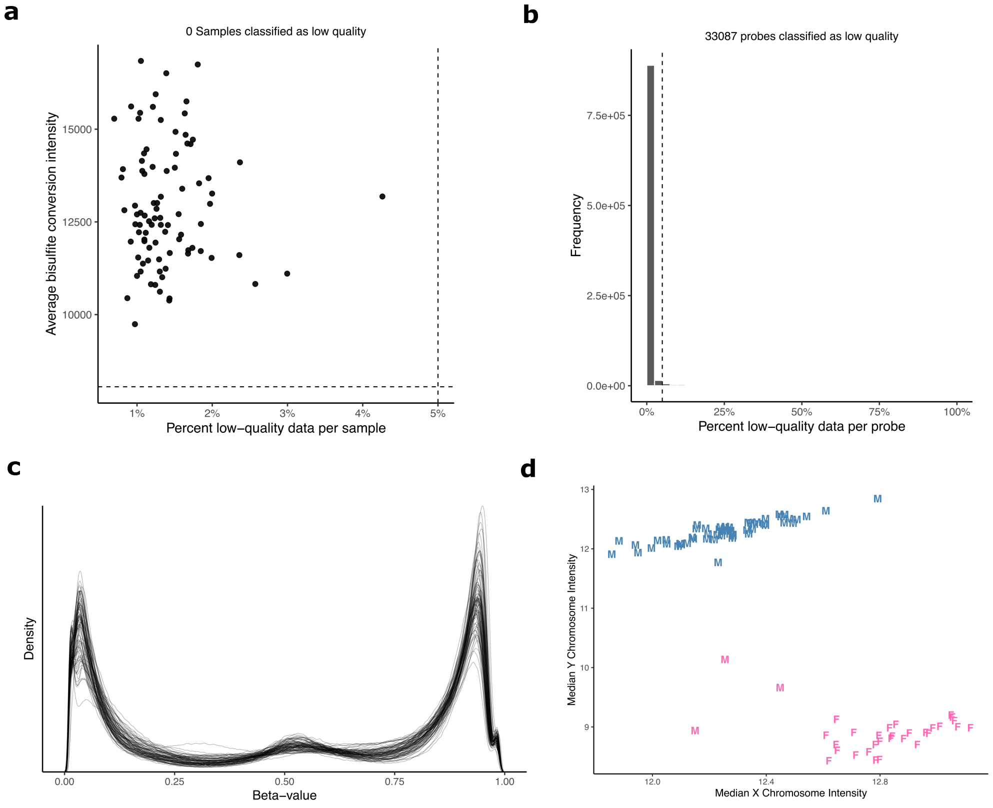


**Supplementary Figure 1: Quality control on methylation data. a** Quality control plot showing percentage of low-quality probes and average bisulfite conversion efficiency per sample. In a given sample, a low quality probe has either a detection p-value $> 10^{-6}$ or less than three beads measured. No samples exceed the exclusion threshold of >5% low quality probes, or exceeding three standard deviations from the population average bisulfite conversion intensity. **b** Per-probe plot, depicting proportions of samples where a given CpG is low quality. Probes deemed low quality in 5% or more of samples, as marked by the right side of the dashed line, were excluded. **c** Beta-value kernel density plot after quality control, filtering, and Normal-Out-Of-Band (NOOB) normalization ($n = 89$). **d** Plot showing sex misclassifications, through median X and Y chromosome intensity. Color corresponds to the methylation-based prediction and the letter (M or F) corresponds to the sample label. 3 samples labeled Male were predicted Female, and 0 labeled Female were predicted male. Maintaining a normal X chromosome intensity while losing Y chromosome intensity may be attributed to Y chromosome loss.

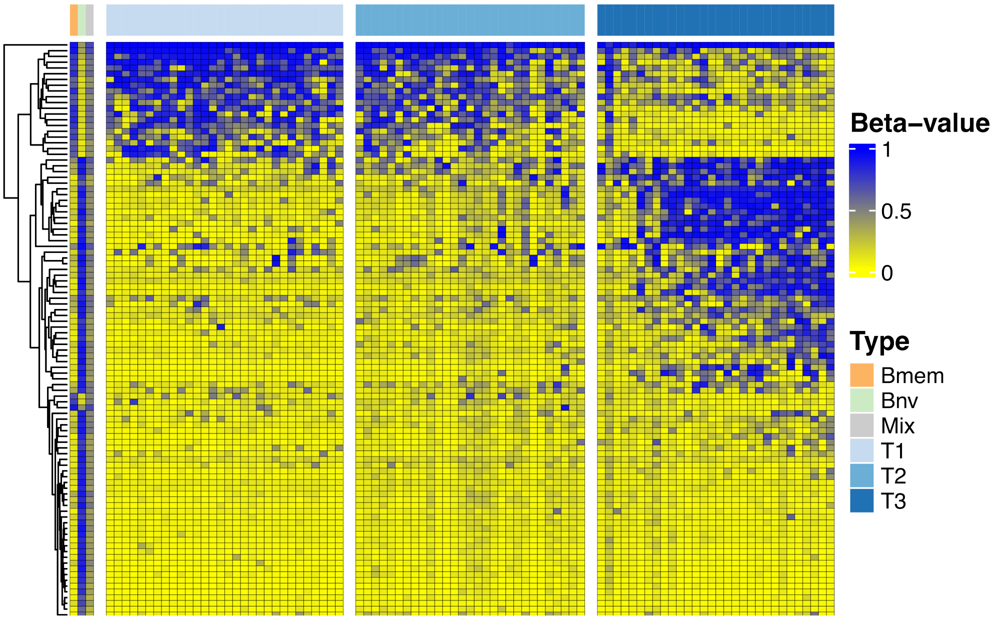


**Supplementary Figure 2: Heatmap of CLL samples at B-memory and B-naive differentially methylated CpGs.** Heatmap shows beta values of the 89 CLL samples, in order of decreasing B-memory proportion (the same order as Figure 1A), at the 100 most differentially methylated CpGs between B-naive and B-memory cells in the reference library. The right-most tertile is also shown in Figure 1C. As in Figure 1C, the pure B-memory and B-naive references are shown, as well as a 60:40 in-silico mixture of B-memory and B-naive references.

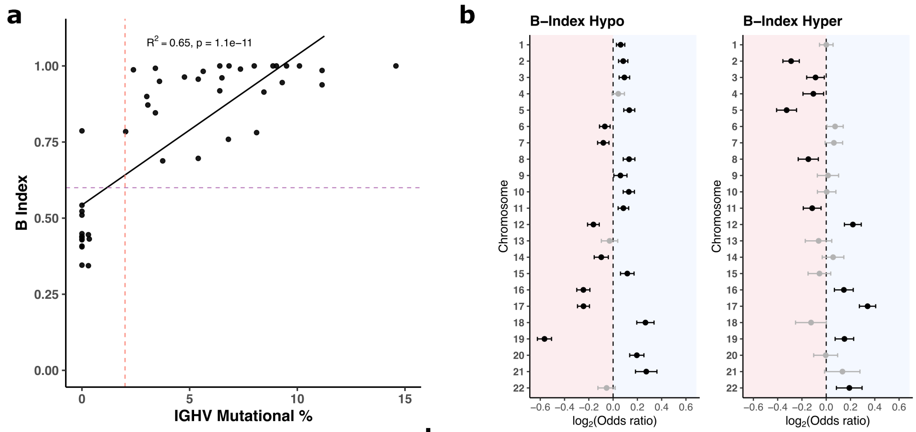


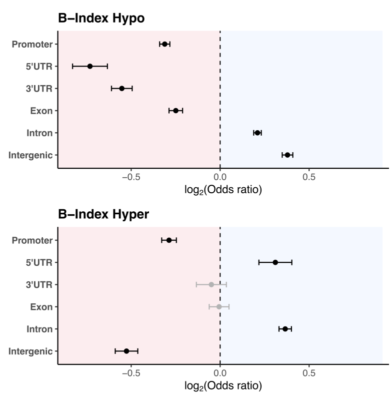


**d**

| Methylation Direction | GO Term | P-value | FDR (BH) |
|---|---|---|---|
| Hypo | cell periphery | 6.83e-48 | 1.52e-43 |
| Hypo | plasma membrane | 3.61e-39 | 4.02e-35 |
| Hypo | signaling receptor activity | 3.59e-20 | 2.00e-16 |
| Hyper | regulation of developmental process | 1.76e-07 | 3.91e-03 |
| Hyper | plasma membrane | 1.31e-06 | 1.42e-02 |
| Hyper | cell periphery | 2.47e-06 | 1.42e-02 |

**e**

| Methylation Direction | KEGG Pathway | P-value | FDR (BH) |
|---|---|---|---|
| Hypo | Cornified envelope formation | 4.68e-14 | 1.72e-11 |
| Hypo | Cytokine-cytokine receptor interaction | 6.08e-10 | 1.12e-07 |
| Hypo | Cytoskeleton in muscle cells | 1.20e-09 | 1.47e-07 |
| Hyper | Phospholipase D signaling pathway | 3.81e-04 | 1.39e-01 |
| Hyper | Glycerolipid metabolism | 7.59e-04 | 1.39e-01 |
| Hyper | Pathways in cancer | 1.64e-03 | 1.66e-01 |

**Supplementary Figure 3: Validation and epigenetic correlates of B-Index. a** Scatterplot of *IGHV* Mutational % and B-Index among 46 samples where percentage information was available. Horizontal purple dashed line is at optimal classification threshold for B-Index of 0.6. Vertical red dashed line is at 2%, the clinically used threshold for U-CLL and M-CLL classification. Black solid line is line of best fit. **b** Forest plot of chromosomal enrichment of hypo and hypermethylated CpGs from EWAS on B-Index. Odds ratio and 95% CI interval shown, in $log_2$ space. Entries in black are significant (FDR $p < 0.05$). **c** Forest plot of genomic context enrichment of hypo and hypermethylated CpGs from EWAS on B-Index, adjusting for CpG island relation using Cochran–Mantel–Haenszel (CMH) test. Odds ratio and 95% CI interval shown, in $log_2$ space. Entries in black are significant (FDR $p < 0.05$). **d** Top 3 hits from two GO pathway enrichment analysis conducted separately on the FDR significantly hypo and hypermethylated CpGs from EWAS on B-Index. **e** Top 3 hits from two KEGG pathway enrichment analysis conducted separately on the FDR significantly hypo and hypermethylated CpGs from EWAS on B-Index.

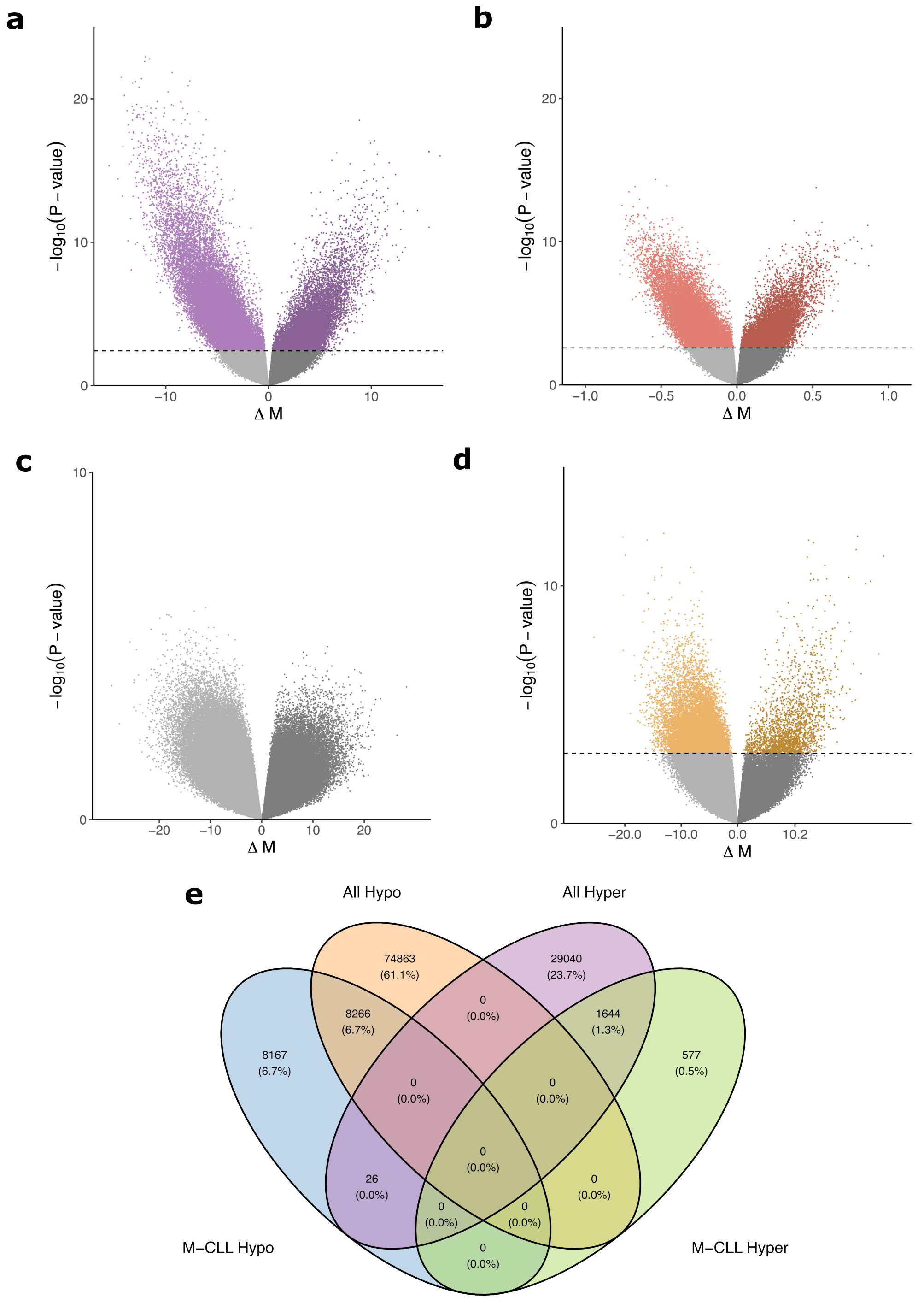


**Supplementary Figure 4: B-Index compared to *IGHV* percentage and epigenetic signal of B-Index within subtypes. a** EWAS of B-Index on cohort of 46 samples where *IGHV* percentage mutation was available, for comparison with (b). 14,776 CpGs were significantly hypermethylated (FDR $p < 0.05$), and 48,982 hypomethylated. **b** EWAS *IGHV* percentage mutation, for comparison with (a). 12,218 CpGs were significantly hypermethylated (FDR $p < 0.05$) and 32,250 hypomethylated. **c** EWAS on B-Index among the 24 U-CLL cases. No FDR-significant hits were observed. **d** EWAS on B-Index among the 56 M-CLL cases. 2,221 CpGs were FDR-significantly hypermethylated, and 16,459 were hypomethylated. **e** Venn diagram of CpGs identified by B-Index EWAS on full cohort, compared with B-Index EWAS within M-CLL subset.

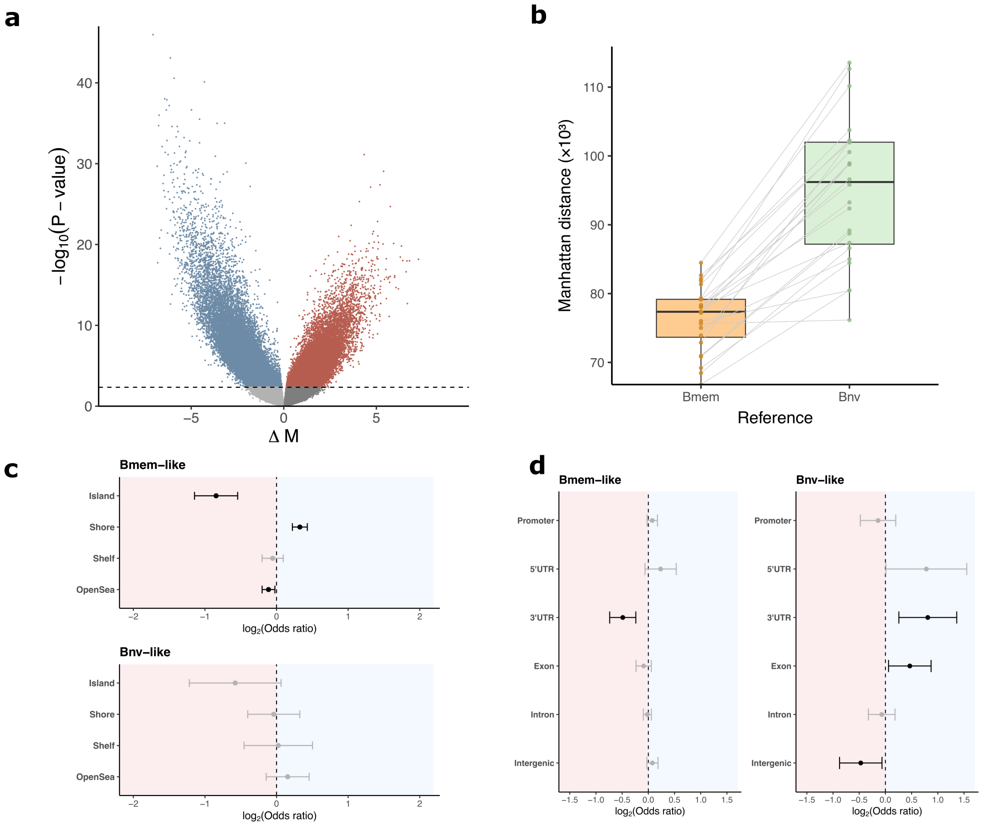


**Supplementary Figure 5: Epigenetic similarities of U-CLL to B-memory and B-naive. a** Volcano plot depicting EWAS between M-CLL and U-CLL. 28,462 were FDR-significantly hypermethylated, and 48,402 hypomethylated. **b** Box plots depicting the Manhattan distance of each U-CLL case to the mean B-memory reference, and mean B-naive reference at the full set of the 682,014 CpGs available across all data. Every U-CLL case has a higher distance from B-naive than B-memory. **c** Forest plot of CpG island context enrichment among CpGs in U-CLL similar (FDR $p < 0.05$) to B-memory and B-naive, adjusting for Illumina probe type using CMH Test. Odds ratio and 95% CI interval shown. Entries in black are significant (FDR $p < 0.05$). **d** Forest plot of genomic context enrichment among CpGs in U-CLL similar (FDR $p < 0.05$) to B-memory and B-naive, adjusting for CpG island relation using CMH test. Odds ratio and 95% CI interval shown. Entries in black are significant (FDR $p < 0.05$).

a

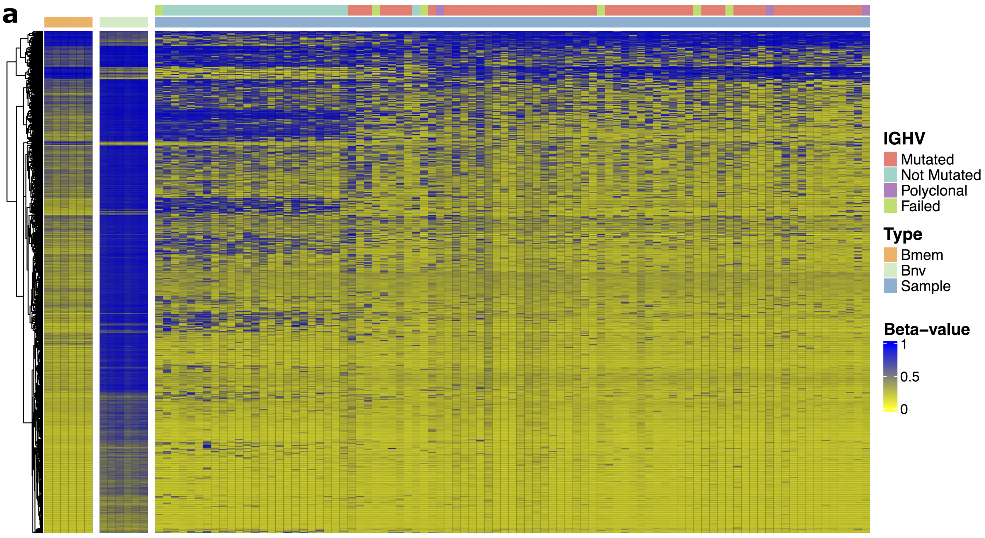


b

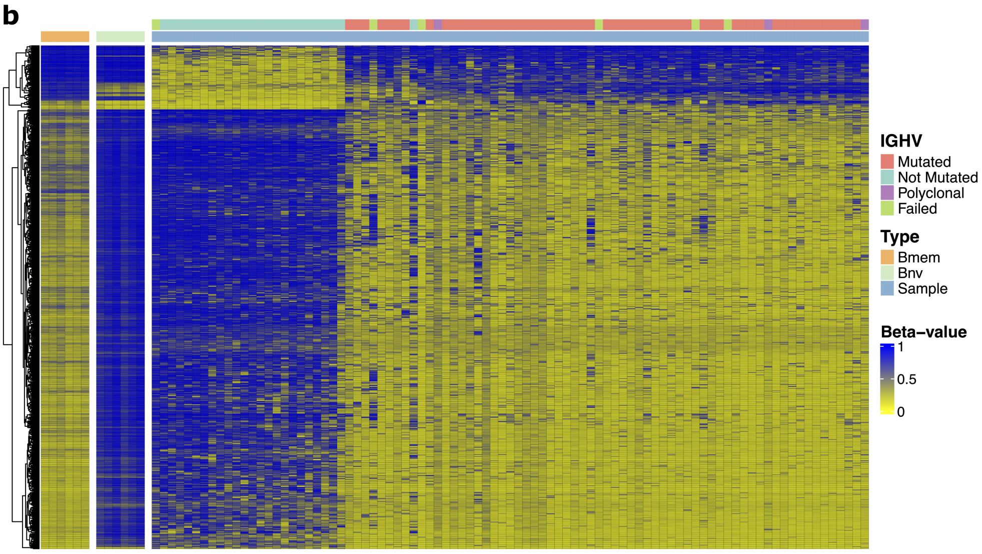


**Supplementary Figure 6: Heatmaps of all CLL samples. a** Heatmap of the top 1000 most significant CpGs from the B-memory v. B-naive EWAS available in our data. B-memory and B-naive references are shown alongside all samples, with IGHV mutational status shown in the top bar, and sorted in order of increasing B-Index from left to right. Heatmap contains the same CpGs from Figure 3B. **b** Heatmap similar to (a), except using top 1000 most CpGS from the U-CLL vs M-CLL EWAS. Heatmap contains the same CpGs from Figure 3C.

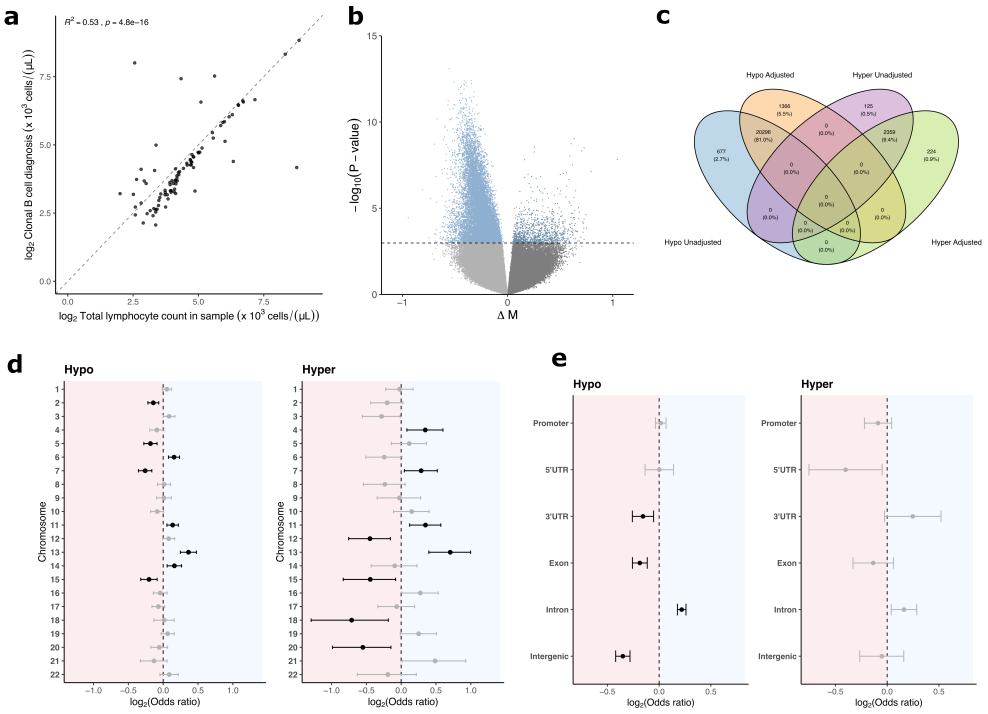


**f**

| Methylation Direction | GO Term | P-value | FDR (BH) |
|---|---|---|---|
| Hypo | cytosol | 1.83e-13 | 4.07e-09 |
| Hypo | lymphocyte activation | 4.67e-13 | 5.20e-09 |
| Hypo | nucleoplasm | 1.27e-12 | 9.41e-09 |
| Hypo | positive regulation of immune response | 6.40e-12 | 2.98e-08 |
| Hyper | structural constituent of chromatin | 8.12e-09 | 1.81e-04 |
| Hyper | chromosome | 3.69e-07 | 3.16e-03 |
| Hyper | nucleosome | 4.26e-07 | 3.16e-03 |
| Hyper | meiotic chromosome segregation | 3.11e-05 | 1.73e-01 |

**Supplementary Figure 7: Epigenetic correlates of tumor burden. a** Scatterplot between the two metrics for tumor burden: total lymphocyte count in sample, and clonal B-cell count at diagnosis (units of thousand cells per microliter). Correlation was conducted in $log_2$ space. $R^2$ was 0.53. **b** Volcano plot depicting EWAS of clonal B-cell count at diagnosis. **c** Venn diagram depicting the number of hypermethylated and hypomethylated FDR $p < 0.05$ significant CpGs from an EWAS on total lymphocyte count, compared to the same EWAS but adjusting for B-Index. **d** Forest plot of chromosomal enrichment among CpGs hypo and hypermethylated with increasing tumor burden. Odds ratio and 95% CI interval shown. Entries in black are significant (FDR $p < 0.05$). **e** Forest plot of genomic context enrichment among significantly (FDR $p < 0.05$) CpGs hypo and hypermethylated with increasing tumor burden, adjusting for CpG island relation using Cochran–Mantel–Haenszel (CMH) test. Odds ratio and 95% CI interval shown. Entries in black are significant (FDR $p < 0.05$). **f** Top 4 hits from gene ontology (GO) analysis on significantly (FDR $p < 0.05$) hypo and hypermethylated CpGs with increasing tumor burden.
